# Genome-wide identification of fitness determinants in the *Xanthomonas campestris* bacterial pathogen during early stages of plant infection

**DOI:** 10.1101/2022.02.07.479439

**Authors:** Julien S. Luneau, Maël Baudin, Thomas Quiroz-Monnens, Sébastien Carrère, Olivier Bouchez, Marie-Françoise Jardinaud, Carine Gris, Jonas François, Jayashree Ray, Babil Torralba, Matthieu Arlat, Jennifer D. Lewis, Adam M. Deutschbauer, Emmanuelle Lauber, Laurent D. Noël, Alice Boulanger

**Author notes:** Correspondence and LIPME, Université de Toulouse, INRAE, CNRS, Université Paul Sabatier, Castanet-Tolosan, France.

## Abstract

Plant diseases are an important threat to food production. While major pathogenicity determinants required for disease have been extensively studied, less is known on how pathogens thrive during host colonization especially at early infection stages. Here, we used randomly barcoded-transposon insertion site sequencing (RB-TnSeq) to perform a genome-wide screen and identify key bacterial fitness determinants of the vascular pathogen *Xanthomonas campestris* pv. *campestris* (*Xcc*) during infection of the cauliflower host plant (*Brassica oleracea*). This high-throughput analysis was conducted in hydathodes, the natural entry site of *Xcc*, in xylem sap and in synthetic media. *Xcc* did not face a strong bottleneck during hydathode infection. 183 genes important for fitness were identified in plant-associated environments with functional enrichment in genes involved in metabolism when only few genes known to be involved in virulence were found to be affected. The biological relevance of 13 genes was independently confirmed by phenotyping single mutants. Notably, we show that the XC_3388, a protein with no known function (DUF1631), plays a key role in the adaptation and virulence of *Xcc* possibly through c-di-GMP-mediated regulation. This study thus revealed yet unsuspected social behaviors adopted by *Xcc* individuals when confined inside hydathodes at early infection stages.

## Introduction

Pests and pathogens of crops cause significant losses in yield and quality. The search for new control strategies relies on our understanding of host-pathogen interactions. *Xanthomonas* bacteria cause disease on more than 400 plant species and are responsible for important losses on various economically important crops worldwide, including rice, citrus, cassava, banana or tomato (Büttner and Bonas, 2010). Among them, *Xanthomonas campestris* pv. *campestris* (*Xcc*), the causal agent of the black rot disease, is the major bacterial pathogen of *Brassica* crops such as cauliflower, cabbage, mustard or radish (Vicente and Holub, 2013). Transmissible by seeds, *Xcc* survives as an epiphyte and enters the plant through leaf organs called hydathodes. Hydathodes are nephron-like structures located at the leaf margin and connect with xylem vessels. Hydathodes mediate guttation and the release of excess xylem sap when transpiration is limited (Cerutti et al., 2017; Cerutti et al., 2019). To initiate infection, *Xcc* enters host tissues via the hydathode pores. After a six-day long biotrophic phase, *Xcc* switches to necrotrophic behavior and destroys the inner epithemal tissue of hydathodes (Cerutti et al., 2017; Luneau et al., 2022a). Simultaneously, *Xcc* accesses xylem vessels and progressively spreads systemically in the plant.

*Xcc* deploys a large arsenal of virulence factors to successfully infect the host plant and complete its life cycle, including two protein secretion systems that are essential for pathogenicity: the type II secretion system (T2SS) which exports enzymes such as plant cell wall degrading enzymes to the extracellular space and the type III secretion system (T3SS), which translocates type III effector (T3E) proteins into the host cells to suppress immune responses and hijack host metabolism (Büttner and Bonas, 2010; Tang et al., 2020). In addition, *Xcc* produces exopolysaccharides (EPS) named xanthan and lipopolysaccharides that protect bacterial cells against environmental stresses and support biofilm formation. Quorum sensing coordinates bacterial behavior, including biofilm dispersal, and is required for disease (An et al., 2020). *Xcc* also possesses a wide range of two-component systems that ensure perception of environmental signals and initiation of adaptive responses (Qian et al., 2008). Altogether, these traits contribute to the fitness of the bacterium during the infection. However, while previous studies have investigated in depth the virulence factors deployed by *Xcc* to implement its pathogenic lifestyle at late stages of infection, very few have looked at the genetic determinants of fitness at early stages of plant infection (An et al., 2020; Timilsina et al., 2020; Luneau et al., 2022a). *Xcc* colonization of hydathodes is associated with sedentary behavior, activation of pathogenicity determinants (T3SS) and expression of high-affinity transporters for nutrient uptake (*e.g*. phosphate, sulfate, nitrate; Luneau et al., 2022). Nevertheless, transcriptome profiling only provides gene expression levels without evaluating their i*n planta* contribution to bacterial fitness.

Transposon insertion mutagenesis coupled with next-generation sequencing technologies now allow the study of bacterial fitness at a genomic scale (Cain et al., 2020; van Opijnen and Levin, 2020). Randomly Barcoded-Transposon insertion site Sequencing (RB-TnSeq) allows the high-throughput evaluation of gene contributions to fitness using a saturated library of bacteria mutagenized by barcoded transposons (Wetmore et al., 2015). Such TnSeq approaches have been widely used to study animal pathogens and have led to the discovery of infectious processes including virulence factors, genes required for transmission between hosts or antibiotic resistance (reviewed in Cain et al., 2020; van Opijnen and Levin, 2020) but have only recently been applied to plant pathogens. To date, TnSeq screens of genes contributing to *in planta* fitness have been performed in *Pantoea stewartii* (on corn; Duong et al., 2018), *Dickeya dadantii* (on chicory; Royet et al., 2019), *Agrobacterium tumefaciens* (on tomato; (Gonzalez-Mula et al., 2019; Torres et al., 2022) and *Pseudomonas syringae* (on bean and pepper; Helmann et al., 2019; Helmann et al., 2020). While these studies provided diverse insights specific to each pathosystem, they all highlighted the importance of metabolic capacities and secretion systems for optimal growth *in planta*.

Here, RB-TnSeq was used to identify the genetic basis of *Xcc* adaptation to the environments it colonizes within the host including hydathodes, xylem sap and synthetic media. We identified essential genes for *Xcc*, as well as genes that contribute specifically to fitness *in planta*. We show that *XC_3388* encoding a hypothetical protein is important for adaptation and pathogenicity of *Xcc* inside cauliflower hydathodes, likely through c-di-GMP regulation, which highlights the importance of social behaviors during plant infection.

## Results

### A saturated library of *Xcc* transposon-insertion mutants identifies 365 essential genes

In order to screen for genes involved in the adaptation of *Xanthomonas campestris* pv. *campestris* strain 8004 to environments encountered during its lifecycle, a library of randomly barcoded insertion mutants was constructed using a *mariner* transposon (Wetmore et al., 2015; Fig. 1A). High-throughput transposon insertion site sequencing allowed the mapping of each insertion site in the *Xcc* genome and their association with specific barcode sequences. After filtering out chimeric reads and non-unique transposon barcodes and computationally excluding insertions located outside the main body of the gene (the central 10% - 90% of the coding sequence length), we identified 192,986 mutants for BarSeq-based fitness analysis (see Methods). The mutant library encompassed 51,853 out of the 77,290 possible *mariner* insertion sites (TA dinucleotides) in the *Xcc* 8004 genome. With a four-fold coverage and one insertion every 100 bp on average, the mutagenesis was saturated and allowed the study of 3,674 out of 4,617 annotated CDS (Luneau et al., 2022a)

**Fig. 1:**
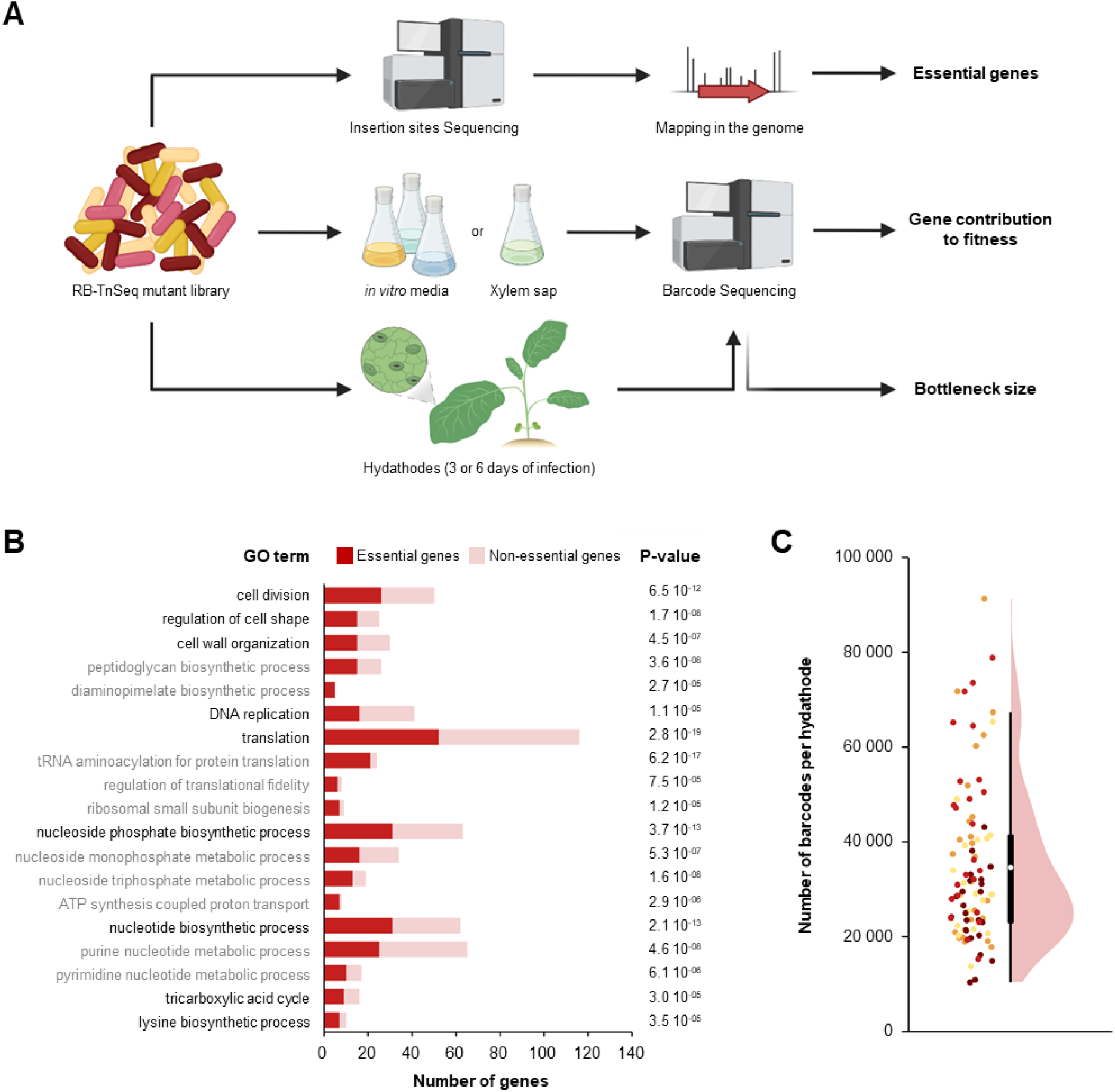
RB-TnSeq approach to identify *Xcc* fitness determinants. **A** Experimental workflow. Barcoded-transposon insertion sites in the *Xcc* 8004::GUS-GFP genome were determined by transposon insertion site sequencing of the RB-TnSeq mutant library and used to identify essential genes. Barcode sequencing of the bacterial population was performed to quantify the relative abundance of all mutants before and after growth of the library in synthetic media, xylem sap and hydathodes. Individual gene fitness values were obtained in each condition. Counting the number of distinct barcodes present within individual hydathodes indicates the bottleneck size for hydathode infection. Created with BioRender.com. **B** Summary of Gene Ontology (GO) functional categories enriched among *Xcc* essential (red) genes compared to non essential (pink) genes. Enrichment significance (P-value) was estimated using Fisher’s exact test. **C** Infection bottleneck in hydathodes determined by counting the number of distinct barcodes found within single cauliflower hydathodes (n=96) three days after dip inoculation with the RB-TnSeq library. Each dot represents a single hydathode. The color represents the four biological repetitions. The boxplot (black) and densityplot (pink) show the distribution of barcode counts across all samples. The white dot indicates the mean value.

Among those 4,617 *Xcc* genes, 80 do not have a TA site in their sequence and cannot be disrupted by a *mariner* transposon. 365 genes were identified as likely essential for viability in *Xcc* strain 8004 based on the presence of TA sites localized in the body of the genes and the absence or very low abundance of mapped transposon insertions (see Methods; Table S1). A Gene Ontology (GO) enrichment analysis highlighted that core cellular functions related to DNA replication, translation, cell envelope, cell division and nucleotide metabolism were most significantly enriched among essential genes (Fig. 1B). The lysine biosynthesis pathway was the only essential amino acid biosynthetic pathway for *Xcc* growth in the rich medium used to construct the library. These results alone significantly improve functional annotation of the genome of the *Xcc* strain 8004 and highlight potential targets for disease control.

### Hydathodes pores do not impose a biologically-significant infection bottleneck for *Xcc*

Prior to any TnSeq analysis, it is important to ensure that a significant part of the mutant library can enter host tissues to minimize sampling artifacts. We thus determined the inoculation bottleneck in *Brassica oleracea* var *botrytis* (cauliflower), the host of isolation of strain 8004 (Turner et al., 1984). Robust hydathode infection was achieved by dip inoculation in the suspension of the mutant library as described (Cerutti et al., 2017) and number of distinct barcodes present in each hydathodes was determined by BarSeq. A mean of 34,582 distinct bacterial mutants were found per hydathode (Fig. 1C) giving a low-end estimate of the infection bottleneck. This result indicates that, in our experimental setup, the entry into hydathodes is not limiting infection because few *Xcc* cells per hydathode are sufficient to initiate infection (Robeson et al., 1989).

Previous work had demonstrated that *Xcc* infection undergoes a biotrophic stage for the first three days before turning to a necrotrophic stage with complete destruction of epithem tissue observable after six days of colonization (Cerutti et al., 2017). We therefore chose to study the fitness of *Xcc* mutants after 3 and 6 days of hydathode colonization, to decouple early infection processes in the hydathodes and adaptive responses in a settled population. Unfortunately, gene fitness values at 3 dpi were noisy and not statistically robust (Fig. S1), likely because there were few generations (around 3 generations total). However, we could still identify 172 genes as directly contributing to *Xcc* fitness in hydathodes (Table S2B and S2C). Almost all the genes found to be important at 3 dpi were also found at 6 dpi (Fig. 2). A GO terms enrichment analysis revealed that functions related to the biosynthesis of cofactors, purine (inosine monophosphate), lipopolysaccharides (LPS) and multiple amino acids were significantly associated with *Xcc* fitness in hydathodes. We also identified phosphate uptake and gluconeogenesis as statistically enriched pathways.

**Fig. 2:**
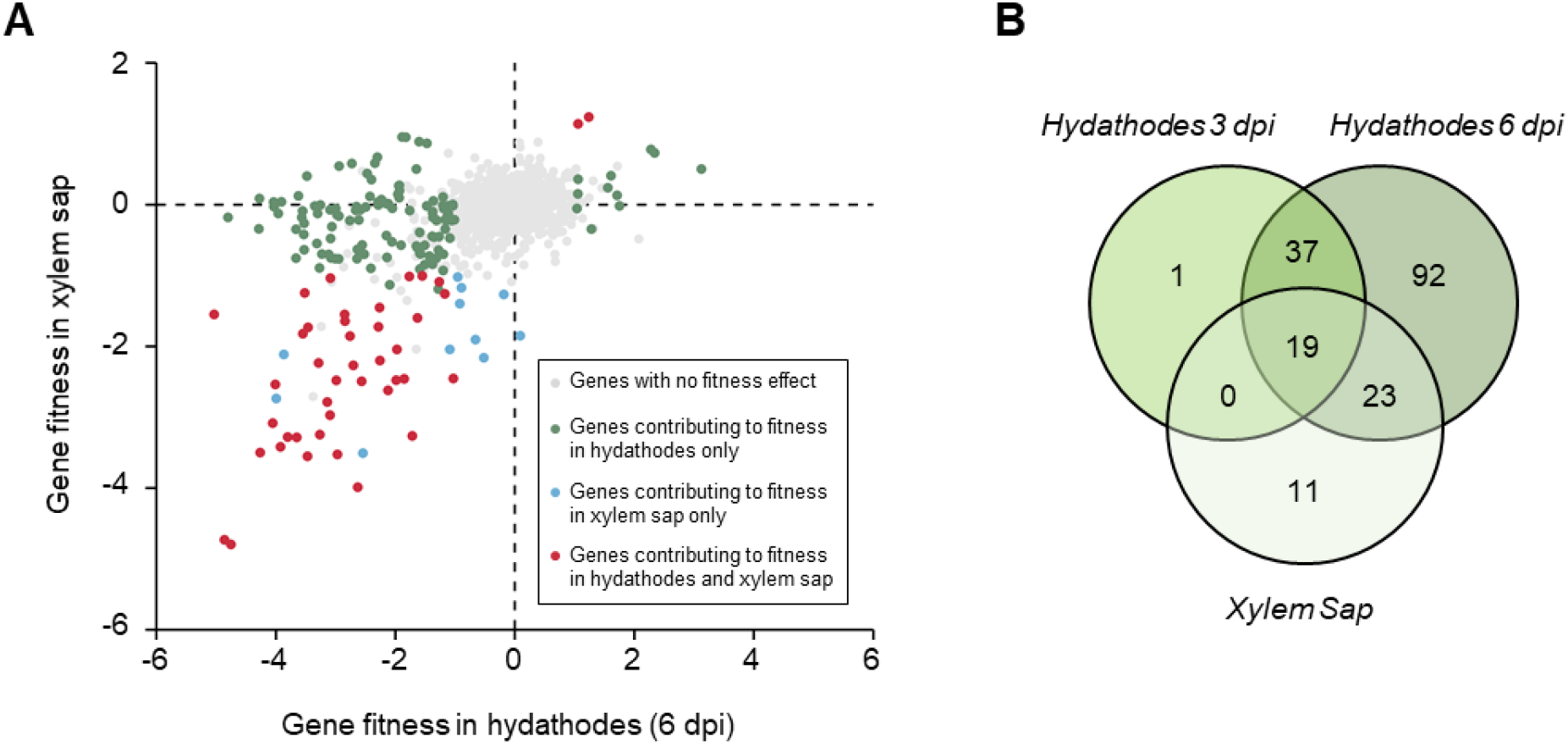
RB-TnSeq screening of *Xcc* fitness determinants during growth in plant-associated environments. **A** Comparison of gene contribution to fitness during growth within hydathodes and in xylem sap. Genes with no fitness effect or that do not fit our robustness requirement are in gray (|f| ≥ 1, |t-score| ≥ 3). Genes found as differentially represented in hydathodes only are in green, in xylem sap only are in blue and in both conditions are in red. Values used here are the mean of 4 biological replicates. **B** Venn Diagram showing the number of genes contributing to *Xcc* fitness during growth in hydathodes at 3 or 6 dpi and in xylem sap.

Comparing the datasets obtained from hydathodes and xylem sap, we identified 19 shared genes for *Xcc* fitness in all three plant-associated conditions investigated here (hydathodes at 3 dpi, 6 dpi and xylem sap; Fig. 2B; Table S2D). We identified genes associated with purine (inosine monophosphate) and vitamin (NADH and Folate) biosynthesis, and TCA cycle-related pathways in all cases. However, these quasi-essential functions are also impacted *in vitro* and are therefore not specific to *in planta* fitness (Fig. S2; Table S2E). Biosynthesis of all amino acids, except cysteine, methionine, proline and valine, was required for optimal multiplication of *Xcc* in hydathodes but was not limiting in xylem sap (Table S2), where only proline biosynthesis was important. Furthermore, among the 53 genes contributing to growth in xylem sap, only 11 genes were specific to the xylem sap condition (Fig. 2B; Table S2). Most of these 11 genes are involved in central metabolic pathways and were also found to be important for growth in artificial media *in vitro* (Table S2). Nevertheless, we detected (p)ppGpp metabolism as specifically important in xylem sap, suggesting that *Xcc* faces nutrient scarce conditions that require a stringent response (Irving et al., 2021; Bai et al., 2022).

### Quantification of the fitness cost of virulence during hydathode infection

Our RB-TnSeq data shows that only a few genes identified as important for *Xcc* fitness in hydathodes are related to virulence functions (He et al., 2007; Fig. S3). Genes involved in EPS biosynthesis and T2SS contribute to *Xcc* fitness, but none of the T3SS machinery components or Type III effectors (T3E), nor quorum sensing signal biosynthesis genes came out of the screen. Yet, we observed a gain of fitness in mutants associated with major regulators of virulence factors, including the *hrpG* and *hrpX* transcriptional activators of the T3SS and the *rpfC* and *rpfG* genes encoding the two-component system involved in quorum sensing signal perception, as well as many transcriptional regulators involved in adaptive responses to the environment such as *xibR* (iron metabolism; Pandey et al., 2016), *ravR* (response to oxygen tension; He et al., 2009), *hpaR* (extracellular proteases production; Wei et al., 2007), *vemR* (motility and EPS production; Tao and He, 2010) and *clp* (EPS and extracellular enzyme production; He et al., 2007b) (Table 1). This result reveals the cross-complementation between individuals associated with the fitness cost of producing virulence factors. It highlights social behavior happening amongst *Xcc* strains in a confined environment such as hydathodes.

**Table 1:**
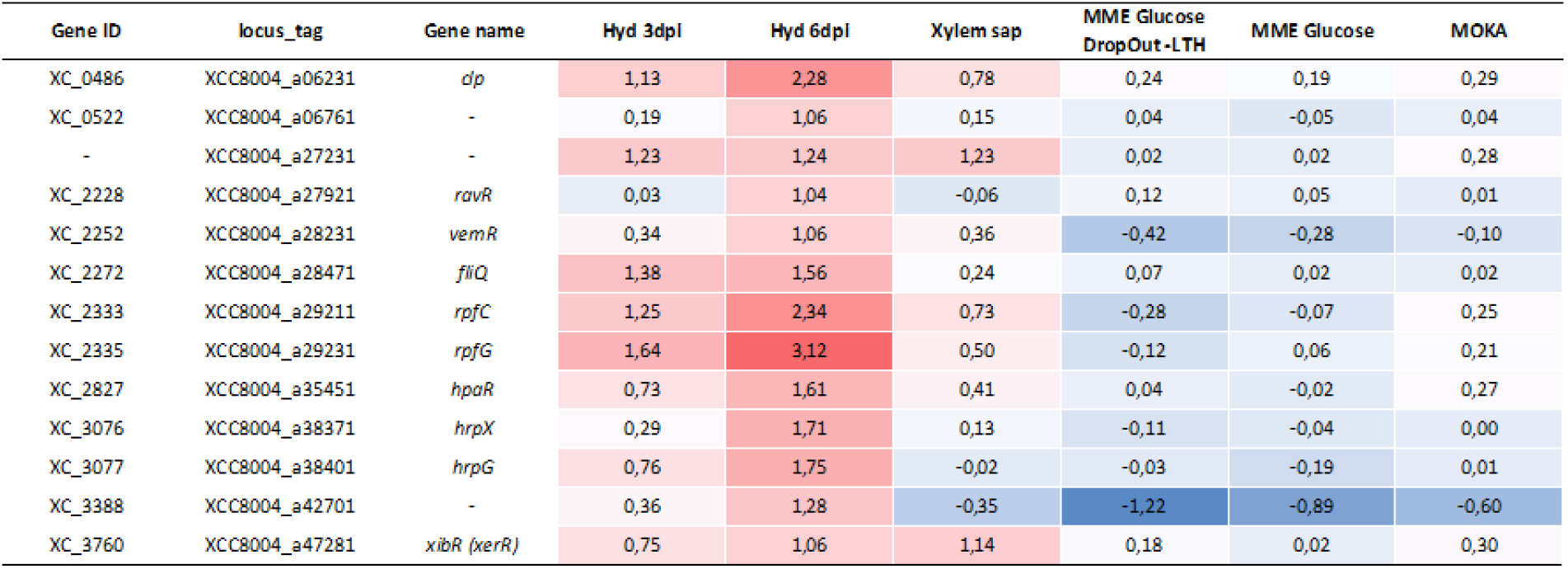
Genes associated with mutants showing a gain of fitness inside hydathodes through RB-TnSeq. Gene fitness values are colored in red when mutants show a gain of fitness and in blue when they show a loss of fitness.

### Contribution of fitness-associated genes to *in planta* multiplication and pathogenicity

To extend the descriptive value of our dataset, we selected 13 candidate genes with either a fitness deficit (9 genes) or a fitness benefit (4 genes) and constructed 12 in-frame deletion mutants (one deletion mutant for both *phaE*-*phcC* genes) for further investigation (Table 2). We assessed the fitness of these 12 mutants by competition assays in dip-inoculated cauliflower hydathodes where each mutant was co-inoculated in a 1:1 ratio against the wild-type strain (Fig. 3). For genes showing a fitness deficit (Fig. 3A), except *XC_1113* and *XC_0555*, deletion mutants showed a deficit in competition, which was complemented. Results obtained with these deletion mutants correlated well with the RB-TnSeq data (Spearman correlation coefficient r = 0.87, p-value = 0.00013; Fig. S4A). This validates the RB-TnSeq screening results and allows the extrapolation of descriptive loss of fitness into the functional importance of mutated genes.

**Table 2:** Candidate genes selected for investigation based on the RB-TnSeq data. Gene fitness values are colored in red when mutants show a gain of fitness and in blue when they show a loss of fitness.

| Gene ID | locus_tag | Predicted Function | Gene name | Hyd 3dpi | Hyd 6dpi | Xylem sap | MME Glucose DropOut-LTH | MME Glucose | MOKA |
| --- | --- | --- | --- | --- | --- | --- | --- | --- | --- |
| XC_0143 | XCC8004_a01841 | 1,4-alpha-glucan branching protein | *glgB1* | -1,93 | -2,66 | -0,08 | -0,08 | -0,15 | -0,08 |
| XC_0449 | XCC8004_a05761 | MarR family transcriptional regulator | - | -1,52 | -2,15 | -0,44 | -0,24 | -0,22 | 0,11 |
| XC_0522 | XCC8004_a06761 | XRE family transcriptional regulator | - | 0,19 | 1,06 | 0,15 | 0,04 | -0,05 | 0,04 |
| XC_0555 | XCC8004_a07161 | hypothetical protein | - | -4,47 | -1,57 | -0,43 | -0,67 | -0,23 | -0,34 |
| XC_1113 | XCC8004_a14101 | TonB-dependent receptor | *bfeA* | -0,46 | -1,94 | 0,10 | -0,02 | -0,05 | 0,09 |
| XC_1431 | XCC8004_a17891 | TetR family transcriptional regulator | - | -0,98 | -2,05 | -0,05 | -0,48 | -0,19 | 0,09 |
| XC_1958 | XCC8004_a24301 | PHA synthase subunit | *phaE* | -1,99 | -2,95 | -0,77 | 0,44 | 0,31 | 0,10 |
| XC_1959 | XCC8004_a24311 | poly-beta-hydroxybutyrate polymerase | *phaC* | -1,79 | -3,27 | -0,90 | 0,43 | 0,22 | 0,03 |
| - | XCC8004_a27231 | stress-induced protein | - | 1,23 | 1,24 | 1,23 | 0,02 | 0,02 | 0,28 |
| XC_2377 | XCC8004_a29681 | imidazole glycerol phosphate synthase subun | *hisH* | -2,08 | -4,04 | 0,04 | -4,30 | 0,18 | 0,20 |
| XC_3076 | XCC8004_a38371 | DNA-binding protein | *hrpX* | 0,29 | 1,71 | 0,13 | -0,11 | -0,04 | 0,00 |
| XC_3253 | XCC8004_a40621 | hypothetical protein | - | -1,62 | -4,05 | -3,08 | -0,63 | -0,16 | 0,05 |
| XC_3388 | XCC8004_a42701 | DUF1631 domain-containing protein | - | 0,36 | 1,28 | -0,35 | -1,22 | -0,89 | -0,60 |

**Fig. 3:**
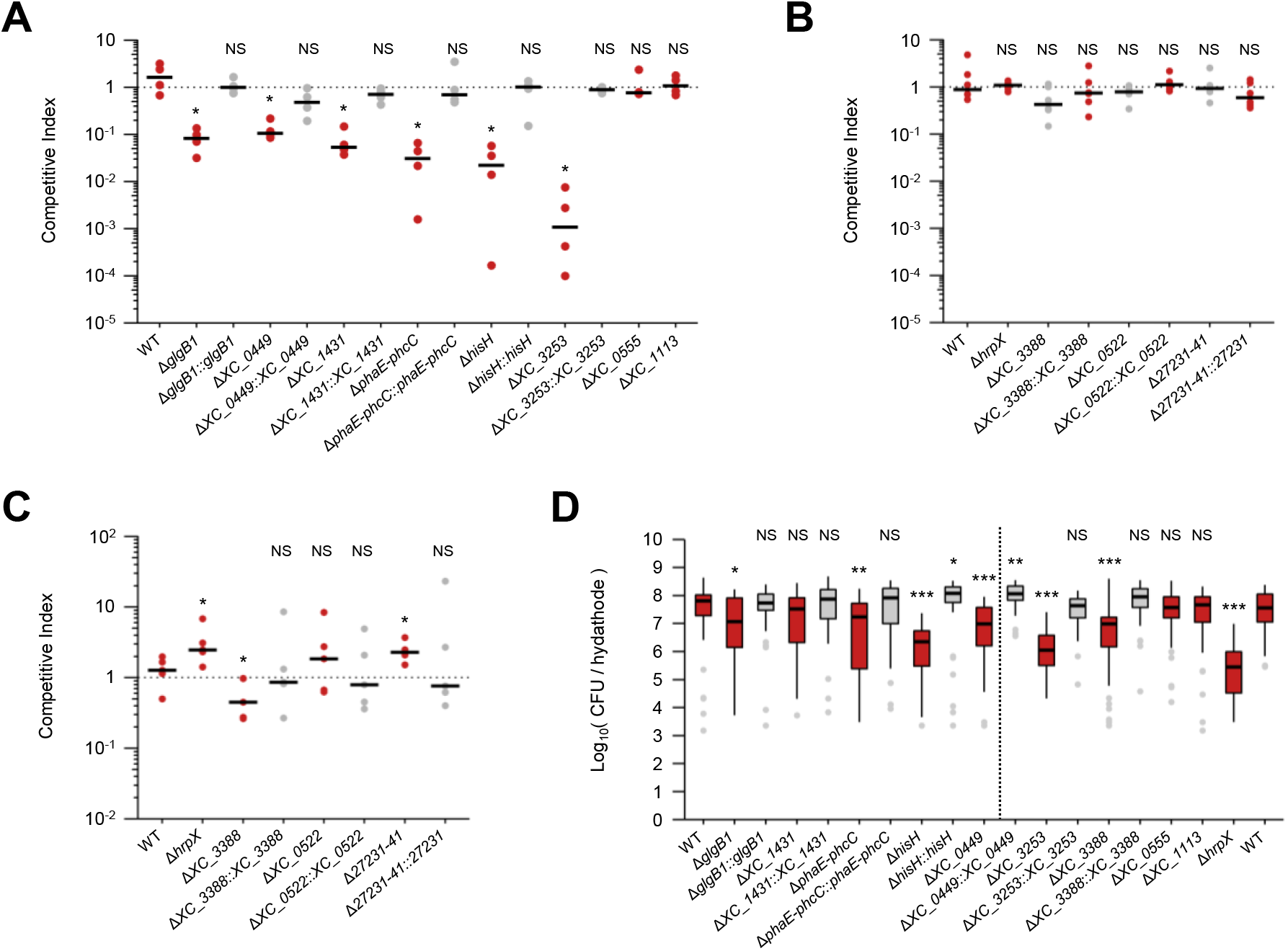
Contribution of candidate genes to *Xcc* fitness during hydathode infection. **A** Competition assays at 1:1 ratio of mutants (red boxes) associated with a loss of fitness in RB-TnSeq or complemented mutants (gray boxes) against the *Xcc* strain 8004∷GUS*-GFP* (hereafter WT*) strain. Dots represent independent biological replicates (N ≥ 4) consisting of 16 pooled hydathodes each. **B** Competition assays at 1:1 ratio of mutants associated with a gain of fitness in RB-TnSeq against the WT* strain. **C** Competition assays at 1:100 ratio of mutants associated with a gain of fitness in RB-TnSeq against the WT* strain. **D** Internal growth curves in individual hydathodes 6 days after dip-inoculation (3 independent biological replicates, n ≥ 37 hydathodes total per strain). Strains separated by the dotted line were inoculated in different experiments. Statistical significance of growth differences between each strain and the WT was assessed with the Wilcoxon test (NS: Not Significant; *: p-value < 0.05; **: p-value < 0.01; ***: p-value < 0.001).

Among the mutants showing a gain of fitness in our RB-TnSeq assay into hydathodes at 6 dpi, we focused on the 3 genes with no known functions (*XC_0522, XC_3388* and *XCC8004_a27231*). Because they showed the same phenotype as mutants associated with known virulence regulators inside hydathodes (e.g. *hrpX* which we used as a control), we hypothesized that these 3 genes could encode novel regulators of adaptation and/or pathogenicity. Competition assays in 1:1 ratio against the wild-type strain did not reproduce the gain of fitness observed in RB-TnSeq (Fig. 3B). However, because mutants in RB-TnSeq assays are present at low frequency in the population, we repeated competition assays with a 1:100 mutant to wild-type ratio (Fig. 3C). It restored the gain of fitness phenotype for the Δ*hrpX* and Δ*27231-41* (knock-out of the overlapping *XCC8004_a27231* and *XCC8004_a27241* genes) mutants showing that these mutants behave like cheaters in a frequency-dependent manner. The failure to restore the gain of fitness for Δ*XC_0522* and Δ*XC_3388* mutants may be due to the fact that the 1:100 ratio isn’t sufficient to reproduce the ratio of these mutants inside our complex library.

Then, we measured the capacity of mutant strains to grow in hydathodes using internal growth curve experiments (IGC; Fig 3D). We observed that all strains tested, except Δ*XC_1431*, show altered growth after six days in hydathodes compared to the WT strain. The discrepancies between CI and IGC (e.g. for Δ*XC_1431*) emphasize that competition assays are better to quantify fine fitness differences but also suggest that co-infection with the WT strain could reveal another mutant phenotype based on cross-complementation.

Finally, among genes with fitness deficit, all except Δ*glgB1* show a defect in pathogenicity when wound-inoculated into cauliflower leaves (Fig. 4A), whereas among genes with fitness benefit, *XC_3388* is the only gene that strongly contributes to disease development (Fig. 4B). These phenotypes were complemented (Fig. 4). Overall, the discrepancy in the phenotypes of *Xcc* mutants highlights that a gene’s contribution to fitness does not always correlate with its contribution to virulence (Fig. S4B).

**Fig. 4:**
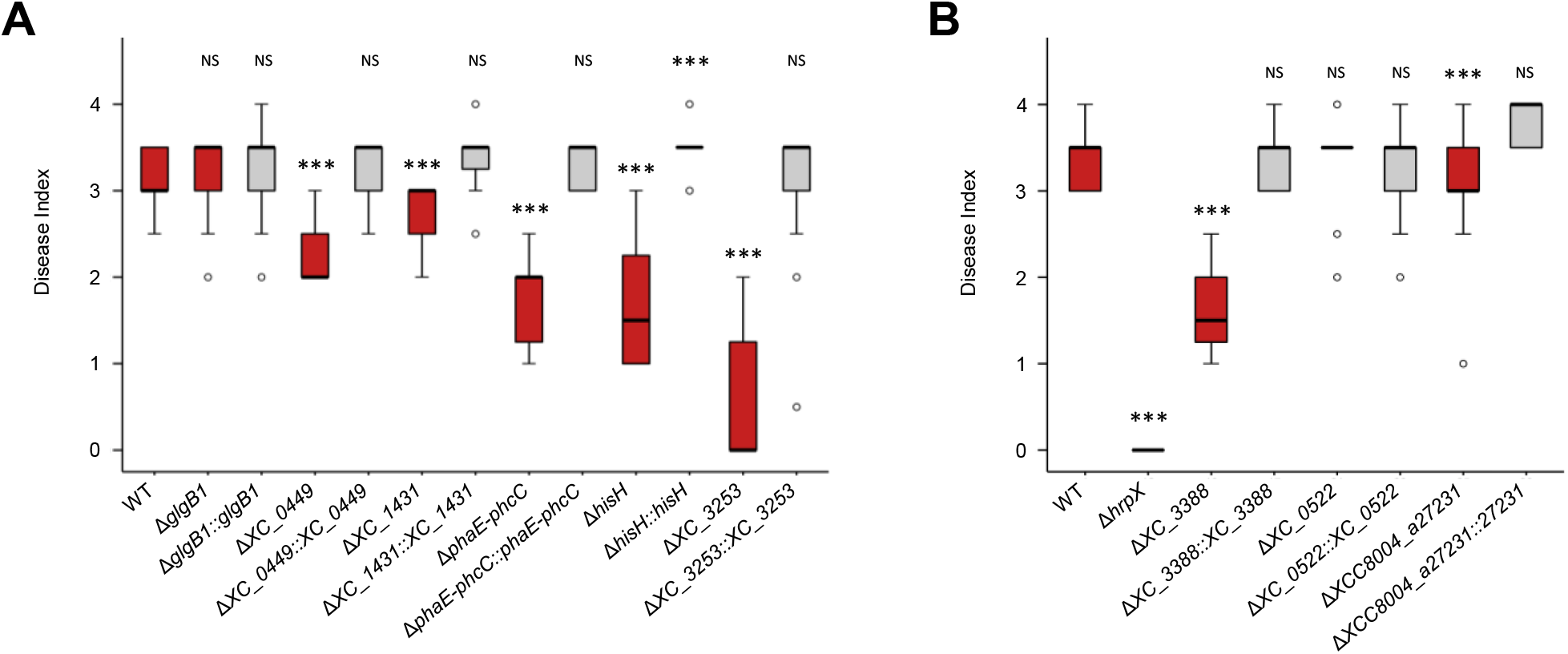
Contribution of candidate genes to *Xcc* pathogenicity. Severity of disease symptoms caused by knock-out (red boxes) and complemented (gray boxes) strains of candidate genes 10 days after piercing inoculation into cauliflower leaf midvein. Strains are grouped based on the phenotype of associated mutants that displayed either a loss (**A**) or a gain (**B**) of fitness in our RB-TnSeq assay at 6 dpi in hydathodes. We determined statistical significance of differences in symptoms severity between each strain and the 8004 WT with the Wilcoxon test (***: p-value < 0.001; NS: Not-significative).

### Investigating proteins of unknown functions involved in pathogenicity

Among our candidate genes, two transcriptional regulators (XC_0449 and XC_1431) and two proteins of unknown function (XC_3253 and XC_3388) were identified as contributing to fitness and disease development during plant infection. To further investigate their potential role in the regulation of the expression of *Xcc* genes, we performed transcriptome analyses of the mutants and complemented strains during growth in MOKA rich medium. Transcriptomic profiling of the Δ*XC_0449* and Δ*XC_1431* strains did not provide any insight into their function or role in *Xcc* physiology (Table S3D and S3E). RNAseq profiling of Δ*XC_3253* showed a deregulation of genes associated with stress responses and motility, which we confirmed by plate motility assays; Fig. S5). Because the *XC_3253* gene is located downstream of an LPS biosynthesis operon, we propose that it is involved in cell envelope formation, though its exact function remains unclear.

We then examined the other gene, XC_3388, which encodes a protein of unknown function. The Δ*XC_3388* mutant displayed pleiotropic physiological alterations. Δ*XC_3388* was hyper-motile on swimming plates, displayed reduced extracellular amylase activity, low EPS production and disturbed biofilm formation (Fig. 5). These observations are consistent with the ‘rough’ colony phenotype visible on agar plates. In addition, metabolic fingerprinting on Biolog plates showed that the Δ*XC_3388* mutant is affected in metabolism of multiple carbon sources including glucose, glutamate, maltose and arbutin (Fig. S6). Evaluation of growth on these carbon sources confirmed the deregulation of metabolic pathways and suggests a role for *XC_3388* in adaptation to environmental conditions.

**Fig. 5:**
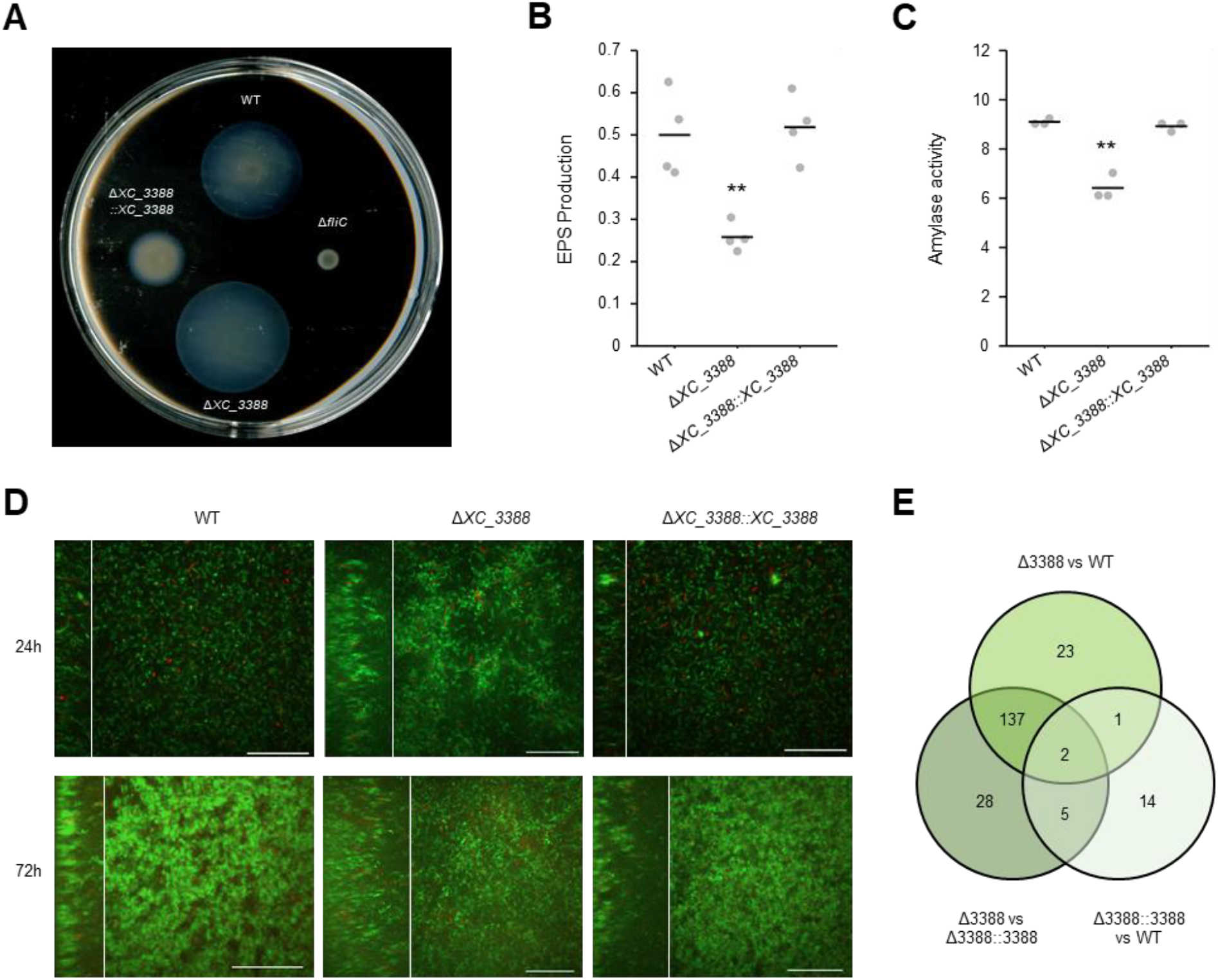
Impact of *XC_3388* on *Xcc* physiology. **A**. Motility assay showing swimming abilities after 48h on soft 0.3% agar plates. The WT strain and the Δ*fliC* non-motile flagellin mutant are positive and negative control of motility, respectively. **B**. Evaluation of exopolysaccharide production after 24h of growth in Moka rich medium. **C**. Evaluation of amylase activity after 24h on Moka plates supplemented with 0.125% potato starch. Statistical significance of differences was assessed with the Wilcoxon test (**: p-value < 0.01). **D**. *Xcc* biofilm architecture. Representative image acquired using spinning disc confocal microscopy after 24h (top panels) and 72h (bottom panels). Cells were grown in chambers with a coverslip containing MME minimal medium supplemented with glucose 20 mM. These observations were performed after addition of propidium iodide to visualize dead cells (red) among live *Xcc* cells (green). Square panels show the biofilm on the *x* and *y* planes, while left rectangles show the biofilm on the *z* plane. Side panel size varies depending on the thickness of the biofilm. Scale bars: 50 µm. **E**. Number of differentially expressed genes during exponential growth in MOKA. Genes were considered differentially expressed if |log_2_FC| ≥ 1.5 and FDR ≤ 0.05.

We then performed a RNAseq transcriptomic analysis comparing the *Xcc* 8004 WT strain with the deletion mutant Δ*XC_3388* and the complemented strain Δ*XC_3388*::*3388* (Fig. 5E and Table S3A). The results confirmed the previous phenotyping assays as deletion of *XC_3388* strongly induces the expression of chemotaxis and flagellar motility genes while repressing EPS biosynthesis genes. We also noted a decrease in expression of ribosomal proteins which can be a response associated with cell stress. Interestingly however, signaling pathways seem significantly deregulated in Δ*XC_3388*. In particular, the expression of many GGDEF domain-containing proteins and diguanylate cyclases, both of which can synthesize c-di-GMP, is induced in the Δ*XC_3388* mutant (Table S3A). This suggests a role of XC_3388 in the metabolism of this important intracellular messenger which could explain the pleiotropic phenotypes observed and supports our hypothesis that XC_3388 could have a regulatory function. The only domain predicted in the sequence of XC_3388 is the domain of unknown function DUF1631 for which no information is available. Structure prediction and comparisons with AlphaFold2 (Jumper et al., 2021) or RoseTTAfold (Baek et al., 2021) could not provide any more insights into XC_3388’s biological role either. Protein sequence analysis by BLAST revealed that *XC_3388* is conserved among the Xanthomonadales and present in some *γ*-proteobacteria genomes. This suggests that the function of this protein is important for these bacteria and that some selection pressure applies here to conserve this function.

## Discussion

### Colonization of hydathodes is constrained by selection rather than bottlenecks

First, by counting the number of barcoded strains entering individual hydathodes, we showed that hydathode pores constitute a very weak bottleneck. Therefore, it seems that hydathodes would be a good entry point for any microbe to colonize the plant. This is consistent with the fact that, unlike stomata, hydathode pores cannot fully close to restrict the passage of bacterial cells (Cerutti et al., 2017). Yet, very few pathogens are known to enter and colonize hydathodes (Cerutti et al., 2019). The paradox posed by the rarity of hydathode-colonizing microbes that infect easily accessible plant tissues suggests that there is strong selection from the plant to prevent hydathode infection. Therefore, it seems that selection rather than stochastic bottlenecking drives the evolution of hydathode pathogens. However, it is unclear what the drivers of this selection are, but could include poor nutrient conditions, or hydathode-specific immune responses. On the other hand, why and how some microorganisms preferentially enter the plant through hydathodes rather than other openings (such as stomata) still remains to be clarified.

### Adaptation to plant environments relies on diverse nutrient sources and biosynthesis pathways

Our screens highlighted the importance of amino acid biosynthetic pathways for *Xcc* fitness in hydathodes and xylem sap. Previous work on cabbage showed that xylem sap contains amino acids, organic acids and sugars in the low millimolar range and that most of these compounds are actively consumed by *Xcc* (Dugé de Bernonville et al., 2014). However, there was no clear correlation between the abundance of amino acids (or their utilization) found by Dugé de Bernonville and colleagues (2014), and gene fitness in xylem sap in our RB-TnSeq study, suggesting that *Xcc* feeds on multiple nutrients in this condition. This hypothesis is confirmed by a recent transcriptomic analysis of *Xcc* during early infection of hydathodes that highlighted the diversity of nutrient scavenging systems expressed in this environment (Luneau et al., 2022a). However, it is interesting to note that the *relA* gene (*XC_1173*) involved in (p)ppGpp biosynthesis was only found to be important for growth in xylem sap. Since accumulation of the (p)ppGpp alarmone triggers the stringent response and allows bacterial cells to face nutrient starvation, it is likely that *Xcc* feeds on anything available in the xylem to ensure multiplication (Bai et al., 2022). In the future, investigating the metabolite composition of guttation fluids and xylem sap will be needed to understand what is available for *Xcc* in the plant. This would complement the current efforts spent on reconstructing bacterial metabolic networks and will ultimately help understand the dynamics of plant infection by phytopathogenic bacteria (Kim et al., 2019; Gerlin et al., 2020). In addition, future studies in leaf mesophyll tissue will enable the generation of a multi-environment map of genetic fitness determinants and thus address questions of tissue specificity.

### RB-TnSeq captures the importance of genes with unknown functions and identifies new pathogenicity determinants

The RB-TnSeq method has gained popularity due to its ability to rapidly perform genome-wide screens and provide phenotypic information about individual gene contributions to bacterial fitness. Price et al. (2018) exploited this feature to investigate the adaptive value of genes from over 30 ecologically distant bacterial strains in a wide variety of conditions. By measuring the contribution of each gene to specific conditions, they could produce co-fitness relationships and associate conditional relevance to 11,000 uncharacterized genes, showing that this method is particularly adapted to unravel the roles of hypothetical proteins. Based on our RB-TnSeq screening *in planta*, we chose to investigate the role of two proteins with unknown function, XC_3253 and XC_3388. In particular, we showed that XC_3388 is pleiotropic and that it contributes to both adaptation to environmental conditions and pathogenicity. While its contribution to pathogenicity has been observed before (Qian et al., 2005), our data allowed us to hypothesize that XC_3388 could be a major regulator of *Xcc* physiology and virulence. The strong deregulation of c-di-GMP-related genes is consistent with the impact on biofilm formation and motility, which are functions known to be regulated by c-di-GMP (Ryan et al., 2007). It is tempting to suggest that XC_3388 could be involved in c-di-GMP homeostasis but the current lack of protein function prediction limits more direct hypotheses. Deeper functional characterization of the protein, particularly of the DUF1631 domain, is therefore needed to shed light on the role of XC_3388 at the molecular and cellular levels.

### Collective behaviors likely increase bacterial fitness during hydathode infection

An important observation of our study is the gain of fitness displayed by mutants of major regulators of pathogenicity during hydathode infection. TnSeq approaches are prone to such counterintuitive results as they evaluate the fitness of individual cells in a population context. In particular, mutations associated with the production of public goods - resources secreted in the extracellular space that are available to the whole population - can be cross-complemented at the population level. Interestingly enough, these mutants in virulence regulators exceed cross-complementation and show an apparent gain of fitness. This strong signal suggests that there is a fitness cost associated with the expression of virulence-related genes during hydathode colonization. Because *Xcc* cells are confined and tightly packed within hydathodes, we suggest (i) that hydathodes constitute a conducive environment for cheater-like behavior and (ii) that RB-TnSeq can finely quantify the cost of virulence during *Xcc* infection.

We showed that cross-complementation occurs with mutants of the *hrpX* transcriptional activator of the T3SS during hydathode infection. These results are similar to what was shown in *Pseudomonas syringae* T3SS-non expressing mutants during tomato leaf mesophyll infection (Rufián et al., 2018). Interestingly, these authors also showed that bi-stable phenotypic heterogeneity emerges in a clonal population and leads to the coexistence of two subpopulations expressing or not-expressing the T3SS (Rufián et al., 2016). Therefore, we suggest that bi-stable T3SS expression is likely to occur in *Xcc* during hydathode colonization. This hypothesis is supported by the frequency-dependent gain of fitness observed in our competition assays. Indeed, the *hrpX* mutant showed a higher fitness relative to the WT strain when inoculated at 1:100 ratio compared to 1:1 ratio. Hydathodes thus seem to be a favorable environment for T3SS-non-expressing cells to grow quickly. Mechanistically, faster growth could be explained by relief of the cost of expressing the T3SS machinery and T3E effectome, all the while benefiting from the immune suppression ensured by cooperators, as it was described for *Salmonella typhimurium* (Sturm et al., 2011). In addition to dividing labor at the population scale, bi-stable expression of the T3SS in *Salmonella* maintains virulence traits fixed in the population by preventing the growth of T3SS-deficient mutants (Diard et al., 2013). From an evolutionary standpoint this is very important because a high abundance of T3SS-deficient mutants could lead to the collapse of infection by *Xcc*. Indeed, according to the Black Queen Hypothesis, emergence of T3SS-deficient mutants would be likely to occur during hydathode infection because it would become evolutionarily more advantageous in the short term to lose these costly genes rather than carry them (Morris et al., 2012). It could allow mutations occurring early in the infection cycle to be fixed in the population and drive the evolution of *Xanthomonas*. This mechanism could explain how non-pathogenic *Xanthomonas* strains lost the T3SS but are still heavily found in the phyllosphere (Vorholt, 2012; Merda et al., 2017). Nevertheless, in the larger context of *Xcc* life cycle, losing the ability to repress host immunity appears to be a weakness since it would limit the infection of the next host and is likely counter-selected.

## Methods

### Bacterial strains and culture conditions

The *Xcc* 8004∷GUS-GFP strain (Cerutti et al., 2017) was used as a recipient for the RB-TnSeq library and all deletion mutants (Table S5). For competition assays, we used the *Xcc* 8004::GUS*-GFP* variant which contains a point mutation inactivating the catalytic sites of both reporter proteins as described in (Luneau et al., 2022b). We cultivated *Xcc* in MOKA rich medium (4 g.l^-1^ Yeast extract, 8 g.l^-1^ Casamino acids, 1 mM MgSO_4_ and 2 g.l^-1^ K_2_HPO_4_; Blanvillain et al., 2007), MME Glucose poor medium (10.5 g.l^-1^ K_2_HPO_4_, 4.5 g.l^-1^ KH_2_PO_4_, 1 g.l^-1^ (NH_4_)_2_SO_4_, 1 mM MgSO_4_, 0.15% (w/v) Casamino acids, 20 mM Glucose; (Arlat et al., 1991) or MME Glucose complemented with DropOut -Leucine/Tryptophan/Histidine supplements instead of casamino acids (10.5 g.l^-1^ K_2_HPO_4_, 4.5 g.l^-1^ KH_2_PO_4_, 1 g.l^-1^ (NH_4_)_2_SO_4_, 1 mM MgSO_4_, 0.15% (w/v) DropOut-LTH, 20 mM Glucose) at 28°C under agitation at 200 rpm or on MOKA-agar plates.

Deletion mutants were constructed with the SacB double recombination method (Schäfer et al., 1994) using pK18 derivatives (Table S4). Complemented strains were obtained by genomic integration *in trans* of selected genes under the constitutive p*tac* promoter using pK18_CompR3 plasmid derivatives (Luneau et al., 2022a; Table S4). Primers designed for all constructions are listed in Table S4. We used the *Escherichia coli* strain TG1 as carrier for the plasmids and the helper strain pRK2073 to introduce plasmids into *Xcc* by triparental conjugation (Figurski and Helinski, 1979; Ditta et al., 1980). *E. coli* strains were cultivated at 37°C on LB-agar or in LB under shaking at 200 rpm. When appropriate, we used the antibiotics rifampicin (50 µg.ml^-1^), kanamycin (50 µg.ml^-1^), spectinomycin (40 µg.ml^-1^) and the fungicide pimaricin (30 µg.ml^-1^).

### Plant material and xylem sap harvest

*Brassica oleracea* var *botrytis* cv. Clovis F1 cauliflower plants were grown under greenhouse conditions and used 4 weeks after sowing. Plants were inoculated in the second true leaf (second leaf above the cotyledons) and placed in growth chambers (8 h light; 22°C; 70% relative humidity). Xylem sap was collected from decapitated cauliflower stems as described in Dugé de Bernonville et al. (2014).

### Construction of the RB-TnSeq library and transposon insertion site sequencing

The RB-TnSeq barcoded transposon insertion library was constructed by conjugating the *E. coli* APA752 donor library containing the barcoded *mariner* plasmid pKMW3 into *Xcc* 8004::GUS-GFP. The mating was performed overnight at a 1:1 ratio on Nutrient Agar (2 g.l^-1^ Yeast extract, 5 g.l^-1^ Peptone, 5 g.l^-1^ NaCl containing 300 μM diaminopimelic acid). Many separate conjugations were then resuspended as a single pooled mixture and spread on MOKA + 50 µg.ml^-1^ kanamycin plates and incubated 2 days at 28 °C to select mutants. All colonies were resuspended in 120 mL MOKA kanamycin and 40 mL of glycerol 80%, aliquoted and frozen at -80 °C. Genomic DNA from *Xcc* mutant library samples was sequenced and barcodes were mapped as previously described in Wetmore et al. (2015).

### Identification of essential genes

Essential genes were predicted as previously described in Price et al. (2018). Genes that lack insertions in our library are likely to be essential in the condition used to build the library (MOKA rich medium). Briefly, for each protein-coding gene, we computed the total read density in TnSeq (reads/nucleotides across the entire gene) and the density of insertion sites within the central 10–90% of each gene (sites/nucleotides). We did not consider the DNA barcodes in this analysis of essential genes. Genes that could have no insertion by chance because of their short length (375 nucleotides or shorter) were excluded from the study.

### Fitness screening in hydathodes

For each biological replicate, a 2 ml-aliquot of the library was thawed and grown overnight at 28°C with shaking at 200 rpm in MOKA + 50 µg.ml^-1^ kanamycin. The culture was washed twice by serial centrifugations (6,000 x g, 10 min) and resuspended in 1 mM MgCl_2_. After measuring the OD_600_, we pelleted 10^9^ *Xcc* cells by centrifugation at 13,000 x g for 5 min and stored them at -80°C to be used as T_0_ samples. Hydathode inoculation was achieved using a protocol adapted from Cerutti et al. (2017). Briefly, cauliflower leaves were dipped for 15 s inside the *Xcc* suspension adjusted to OD_600_ = 0.1 (10^8^ cfu/ml) in 1 mM MgCl_2_ + 0.5% v/v Tween 80. Plants were then placed in growth chamber conditions and covered for 24 h with a mini-greenhouse lid to reach 100% relative humidity. The lid was removed and the infection was allowed to proceed for 3 or 6 days. *Xcc* populations were retrieved by macro-dissection of hydathodes (16 per leaf) using a 1.5 mm-diameter Biopsy Punch (Miltex®, Electron Microscopy Sciences) and bead-grinding in 1 mM MgCl_2_ for 2 min at 30 Hz. In total, we collected around 1,650 hydathodes for each of the four biological replicates. DNA was extracted using the Promega Wizard® Genomic DNA Purification Kit in which we added a protein degradation step by incubating the samples with 400 µg/ml of proteinase K at 37°C for 1h before protein precipitation. Samples were stored at -80°C until preparation for barcode sequencing.

### Bottleneck assays in hydathodes

Hydathode inoculation and sampling were performed as described before (*Fitness screening in hydathodes* section). After three days in the plant, 8 hydathodes per leaf were retrieved and bead-grinded in 200µl of 1mM MgCl_2_ for 2 min at 30 Hz. From this suspension, 10 µl were used to perform serial dilution, plated on MOKA-agar + kanamycin 50 µg.ml^-1^ and counted to check that each hydathode was well infected and contained the same amount of bacterial cells [around 10^5^ CFU/hydathode]. We performed 4 biological replicas of 3 plants each for a total of 96 hydathodes. Individual hydathode homogenates were spotted on sterile filters, placed on MOKA-agar + kanamycin 50 µg.ml^-1^ and incubated three days at 28°C. Grown populations were individually resuspended by mixing filters with 1.5 ml of MOKA and pelleted (6,000 x g, 10 min). DNA was extracted from each pellet and used for barcode sequencing as described before (*Fitness screening in hydathodes* section).

### Barcode sequencing and calculation of gene fitness

Barcode sequencing, mapping, and analysis to calculate the relative abundance of barcodes were done using the RB-TnSeq methodology and computational pipeline developed by Wetmore et al. (2015); the code is available at https://bitbucket.org/berkeleylab/feba/. For each experiment, fitness values (|f|) for each gene were calculated as a log2 ratio of relative barcode abundance after library growth in a given condition divided by relative abundance in the time0 sample. Fitness values were normalized across the genome so the typical gene has a fitness value of 0.

We considered genes to be important for *Xcc* fitness when the mutants were strongly affected, *i.e*. gene fitness |f| ≥ 1, and when this phenotype was deemed robust, *i.e*. |t-score| ≥ 3 (Wetmore et al., 2015; Helmann et al., 2019). We retained genes that match those criteria in at least three of the four independent biological replicates. Enrichment analysis considering gene ontology was conducted with R (version 4.0.4) using the topGO package version 2.40.0 using Fisher’s exact test and p-value < 0.01 (Alexa et al., 2006).

### Competition assays

We mixed the *Xcc* strain 8004∷GUS*-GFP* (hereafter WT*) and deletion mutants in the 8004∷GUS-GFP background to assess the fitness of the latter as described in (Luneau et al., 2022b). Briefly, we inoculated cauliflower hydathodes with the same protocol as for the RB-TnSeq assay except that we adjusted the inoculum suspensions to OD_600_ = 0.05 for each strain (final OD_600_ = 0.1) in 1mM MgCl_2_ + 0.5% v/v Tween 80. In order to maximize the throughput of the validation assays, we chose to inoculate only one plant for each of the mixtures. We plated inoculation suspensions to use as T_0_ reference samples as well as 6 dpi samples consisting of 16 pooled and ground hydathodes. The Competitive Index (CI) of each strain was calculated as the ratio of the relative frequencies of mutant vs. reference strains in the 6 dpi infected sample compared to the T_0_ inoculum (Taylor et al., 1987; Macho et al., 2007). Hence, we consider a CI > 1 as a gain of fitness of the mutant compared to the WT* reference strain and a CI < 1 as a loss of fitness. We performed at least 4 independent biological replicates of the competition assays for all strains tested.

### Pathogenicity assays

We evaluated bacterial aggressiveness of the mutant strains by piercing-inoculation in the midrib of cauliflower leaves using a needle dipped in a *Xcc* inoculum adjusted at OD_600_ = 0.1. We assessed disease progression at 10 days post-inoculation according to the annotation scale presented in (Luneau et al., 2022a) in three independent biological replicates of five individual plants each.

### Internal Growth Curves (IGC) in hydathodes

Bacterial populations in hydathodes were determined after dip-inoculation as described above. After six days, hydathodes were sampled using a cork borer (diameter, 1.5 mm), individually grind using a Tissue Lyser MM 400 grinder (Retsch), two times 30 seconds at a frequency of 30/sec, in 1.2 mL deepwell plates containing 5-6 glass beads (2 mm diameter) per well and 200 µL of sterile water. The homogenates were serially diluted in sterile water and 5 µl drops were spotted three times on MOKA plates supplemented with rifampicin and pimaricin. Plates were incubated at 28°C for 48h and colonies were enumerated in spots containing 1 to 30 colonies. Bacterial densities in leaves were calculated as log CFU/cm^2^.

### Phenotyping assays

Motility, exopolysaccharide production and protease activity were assessed as described in Luneau et al. (2022a). For amylase activity assays, 5 μL of an overnight culture adjusted to 4.10^8^ cfu/mL were spotted on plates containing 15 mL of MOKA agar with 0,125 % potato starch (prolabo, VWR) supplemented with 30 µg/mL pimaricin. Plates were incubated at 28 °C and imaged 24 hours post inoculation. Diameters of colonies and halos of degradation were measured by washing plates using distilled water and stained using lugol solution until a strong blue coloration and starch degradation halo appeared. Calculation of enzymatic activity was performed as follows: PrA = [π(r_halo_)^2^-π(r_colo_)^2^]/ [π(r_colo_)^2^] with “r” being the radius. Each experiment was biologically replicated at least three times.

For biofilm visualization, strains were grown overnight in MOKA rich medium, washed two times with sterile water before resuspended in MME minimal medium supplemented with 20 mM glucose at a final concentration of 10^8^ cells/ml. Five ml of each strain suspension were distributed in 6-well plates (VWR tissue culture plate) containing a sterile borosilicate coverslip in the bottom. One plate per time point is required. Plates were incubated without agitation at room temperature. Before imaging, 3 µl of propidium iodide were added to each well and incubated for 10 minutes. Biofilms were visualized using a spinning disk microscope with a 60x immersion lens.

### Fitness screening *in vitro*

The RB-TnSeq library was grown in MOKA for 4 hours at 28°C with 200 rpm shaking and washed twice in MgCl_2_ 1mM. We pelleted 10^9^ *Xcc* cells by centrifugation at 13,000 x g for 5 min and stored them at -80°C to be used as T_0_ samples. The *Xcc* inoculum was adjusted to OD_600_ = 0.05 (5.10^7^ cfu/ml) in 10 ml of either 0.22 µm-filtrated cauliflower xylem sap, MOKA rich medium, MME poor medium + 20 mM Glucose or MME + 20 mM Glucose + 0.15% DropOut -LTH. We incubated all cultures at 28°C under agitation at 200 rpm. Because the differences in fitness become more noticeable over time, the clarity of RB-TnSeq results depends on the number of generations. Therefore, we allowed bacterial growth for up to 6-7 generations in all media. However, except in MOKA rich medium, *Xcc* growth could not attain this number of generations before reaching the stationary phase. Therefore, we restarted cultures at the end of the exponential phase by centrifugation (6,000 x g, 5 min) and resuspension of the whole population in a larger volume of fresh medium at OD_600_ = 0.05. At the end of growth, we pelleted 10^9^ *Xcc* cells by centrifugation at 13,000 x g for 5 min and stored the pellets at -80°C until DNA extraction (same protocol as before).

### Carbon and nitrogen substrates phenotyping

Utilization of carbon and nitrogen sources by *Xcc* were assayed using the Biolog phenotype microarray plates PM1, PM2A and PM3B (PM; Biolog, Hayward CA) according to the manufacturer’s protocol. *Xcc* cells were grown in rich MOKA medium overnight at 28°C. Cells were collected and resuspended in IF-0 (for PM1 and PM2A) or IF-0 20mM glucose (for PM3B) supplemented with 1× Biolog Redox Dye Mix H, at initial OD_600nm_ = 0.08 for plates inoculation (100 µl/well). Plates were incubated for 72 h in the Omnilog device (Biolog, Hayward, CA) at 28°C. Two independent replicates were performed. Results were analyzed using the Scilab software.

### Growth measurements *in vitro*

*In vitro* growth curves were obtained by growing 200 µL of *Xcc* suspensions in 96-well flat-bottom microtiter plates (Greiner) within a FLUOStar Omega apparatus (BMG Labtech, Offenburg, Germany) at 28°C. After an overnight preculture in MOKA rich medium, cells were harvested by centrifugation at 9 500 x g for 4 minutes, washed and resuspended in MME minimal medium. Bacterial suspensions inoculated at an optical density at 600 nm (OD_600_) of 0.15 were prepared in MME or MME supplemented with either glucose, glutamate, maltose or arbutin at 20 mM and MOKA media. For each experiment, we performed 4 replicates coming from 2 independent growth. The microplates were shaken continuously at 700 rpm using the linear-shaking mode. Each experiment was repeated 3 times from which one representative experiment is shown.

## Acknowledgments

J.S.L. was funded by a PhD grant from the French Ministry of Higher Education, Research and Innovation. J.S.L, M-F.J, S.C, E.L, M.A, L.D.N and A.B were funded by grants from the Agence Nationale de la Recherche XBOX (ANR-19-CE20-JCJC-0014-01). J.S.L was funded by the Fédération de Recherche Agrobiosciences, Interactions et Biodiversité through project CHACCCAL. A.B. and O.B. were funded by the Fédération de Recherche Agrobiosciences, Interactions et Biodiversité and TULIP Labex through project 31000509_FRT4. This study is set within the framework of the “Laboratoires d’Excellences” (LABEX) TULIP (ANR-10-LABX-41) and of the “École Universitaire de Recherche” (EUR) TULIP-GS (ANR-18-EURE-0019). Research in J.D.L.’s lab is supported by USDA ARS Grants 2030-21000-046-00D, 2030-21000-050-00D, and NSF Directorate for Biological Sciences Grant IOS-1557661. All authors benefited from the COST action CA16107 EuroXanth.

---

**Fig. S1:**
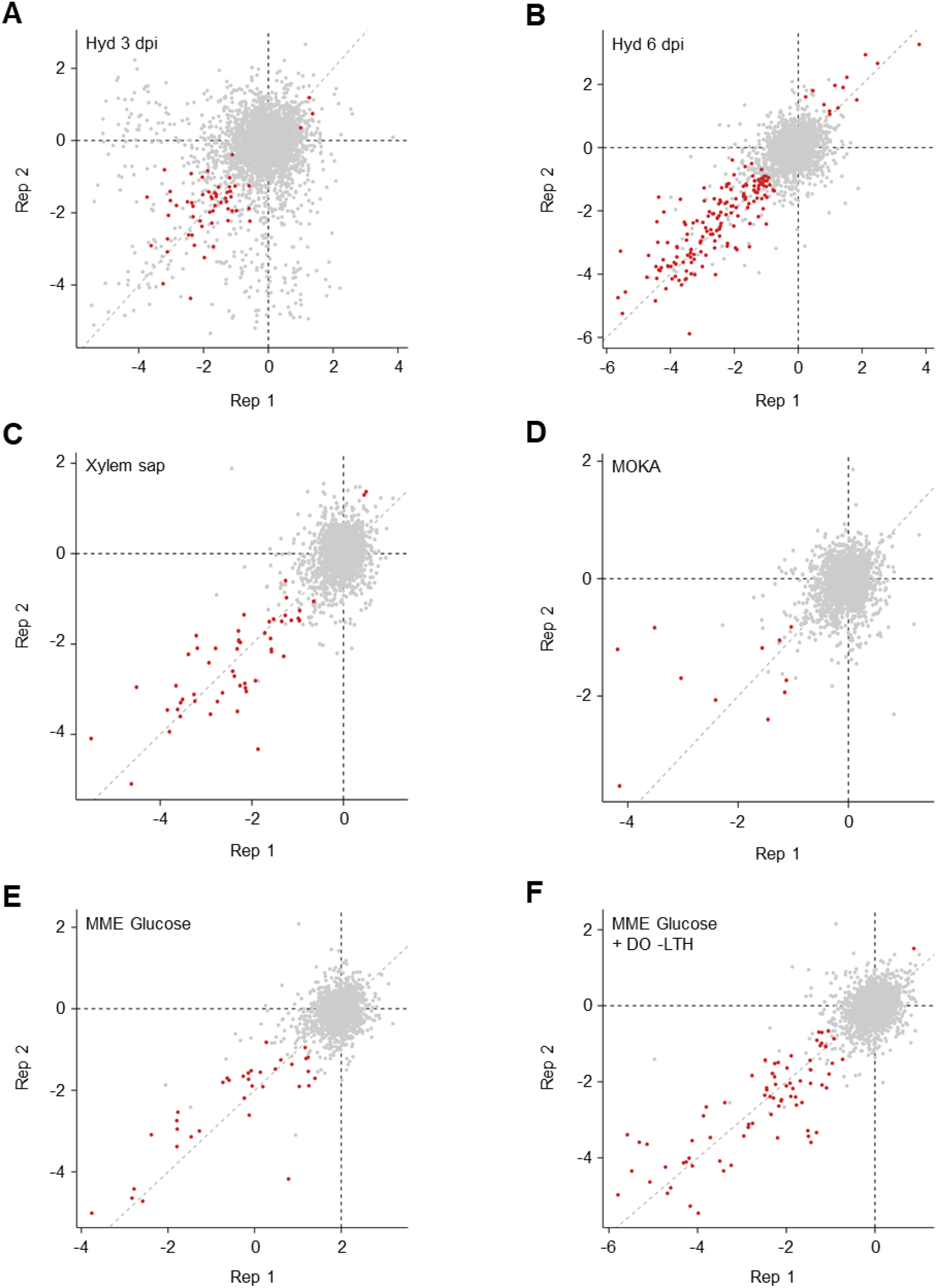
Correlation between biological replicates of RB-TnSeq assays shows experimental reproducibility. We investigated biological replicates correlate for the environments tested in our study. **A** Hydathodes 3 dpi. **B** Hydathodes 6 dpi. **C** Xylem sap. **D** MOKA rich medium. **E** MME Glucose. **F** MME Glucose + DropOut –LTH. Each dot represents a gene. Red dots show the genes selected according to our criteria (|f| > 1 & |t-score| > 3). The grey dashed line traces the y = x ideal correlation between the two replicates. Two out of four biological replicates are shown for each condition and are representative of the pairwise correlations between all the replicates.

**Fig. S2:**
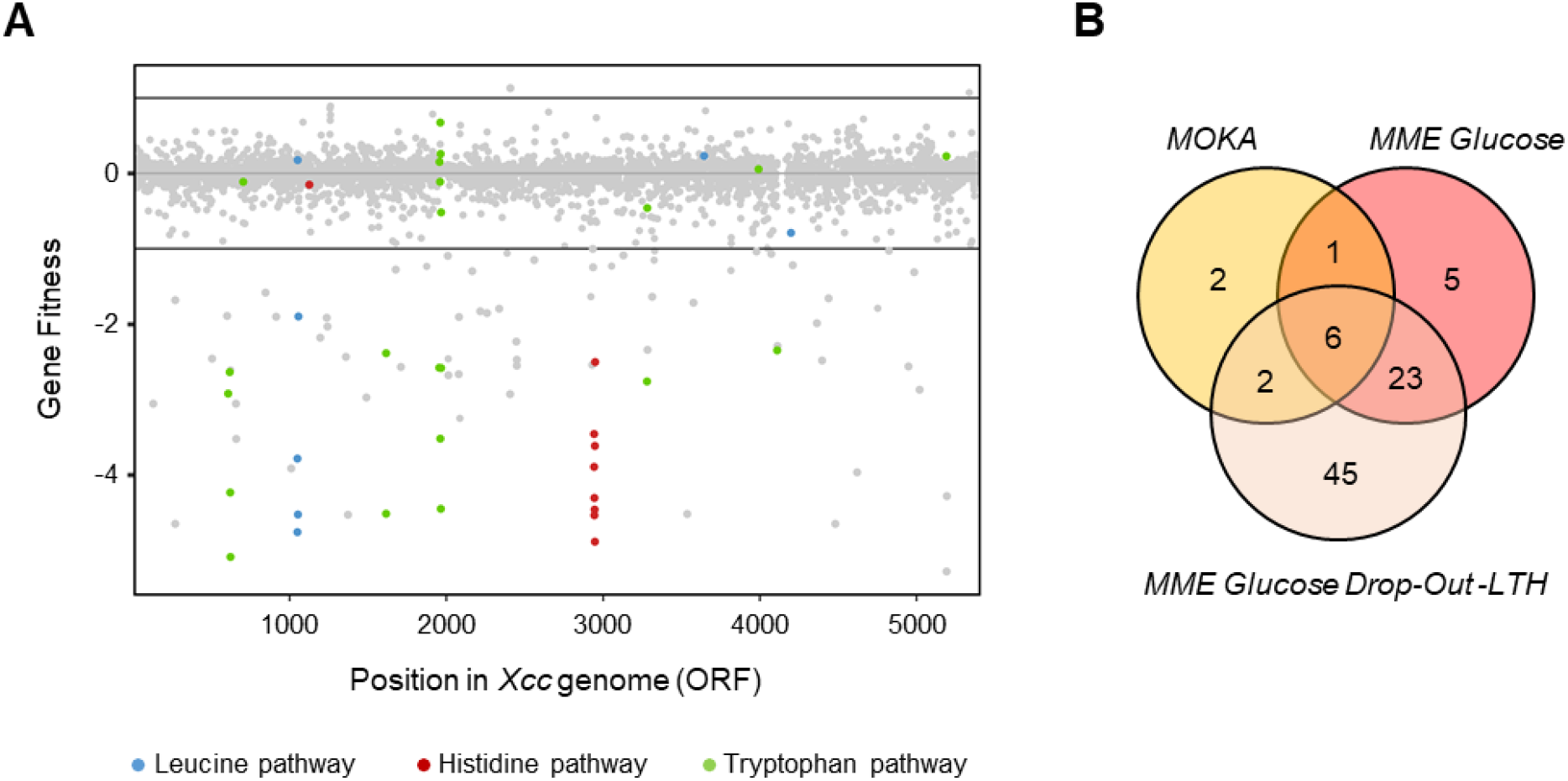
Growth of the RB-TnSeq library *in vitro* identifies genes contributing to multiplication in plant-independent conditions. **A** Genome-wide gene fitness assessment in minimal medium MME Glucose DropOut –LTH medium. Genes related to the biosynthesis of leucine, histidine and tryptophan are highlighted. The expected loss of fitness in these auxotrophic mutants grown in a medium lacking the three amino acids confirms the utility of our RB-TnSeq library to identify conditionally-relevant genes and validates the quality of the readout for evaluating gene fitness. **B** Venn Diagram showing *Xcc* genes contributing to *in vitro* growth which are quasi-essential and not specific to the plant-associated environments tested. The MOKA rich medium was used to construct the library.

**Fig. S3:**
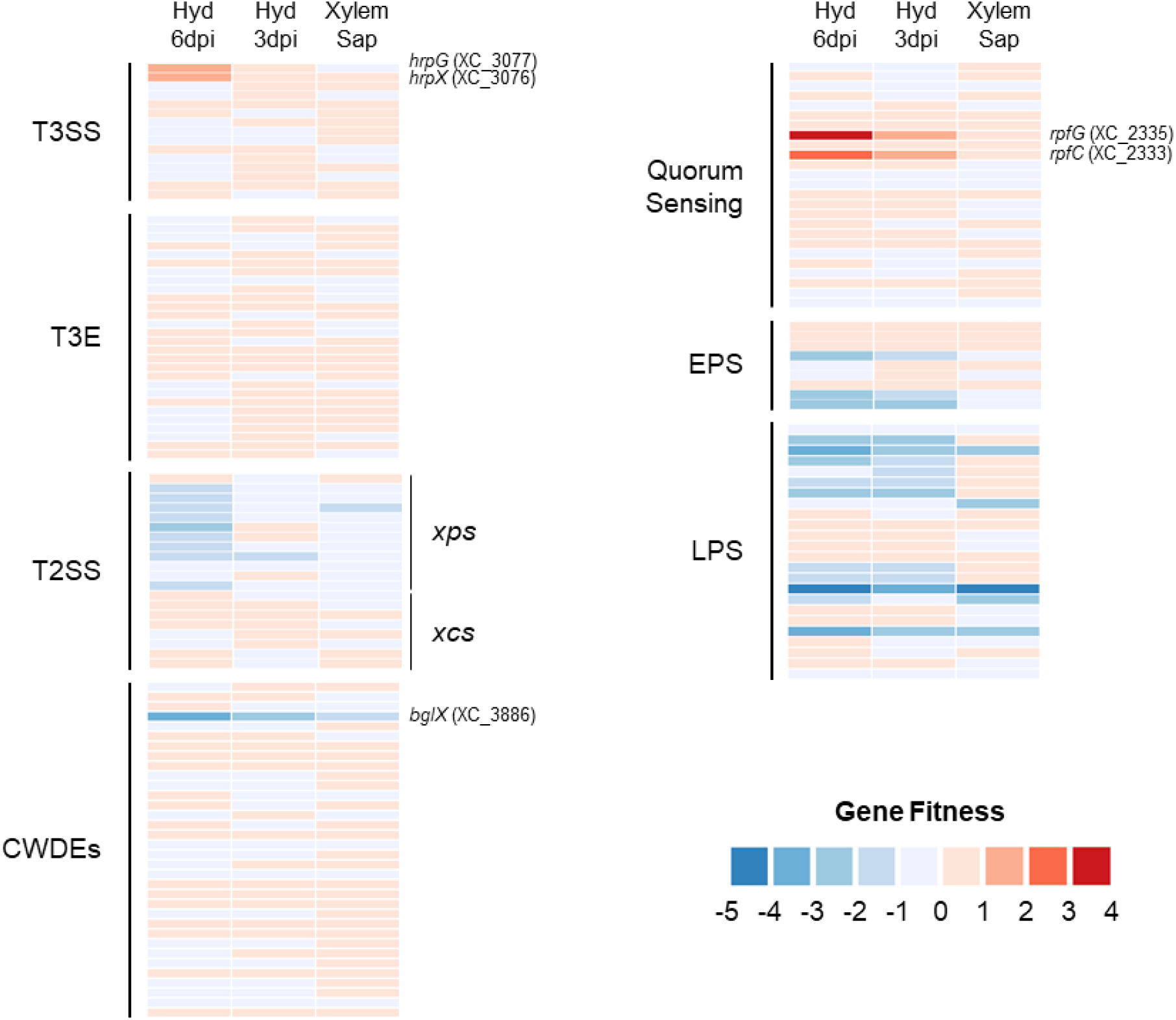
RB-TnSeq quantification of virulence factors contributing to *Xcc* fitness in plant-associated conditions. Fitness profile of *Xcc* genes associated with virulence functions according to He et *al*. (2007). T3SS : Type III Secretion System; T3E : Type III Effector; T2SS : Type II Secretion System; CWDEs : Cell Wall Degradation Enzymes; EPS : Exopolysaccharide biosynthesis; LPS : Lipopolysaccharide biosynthesis.

**Fig. S4:**
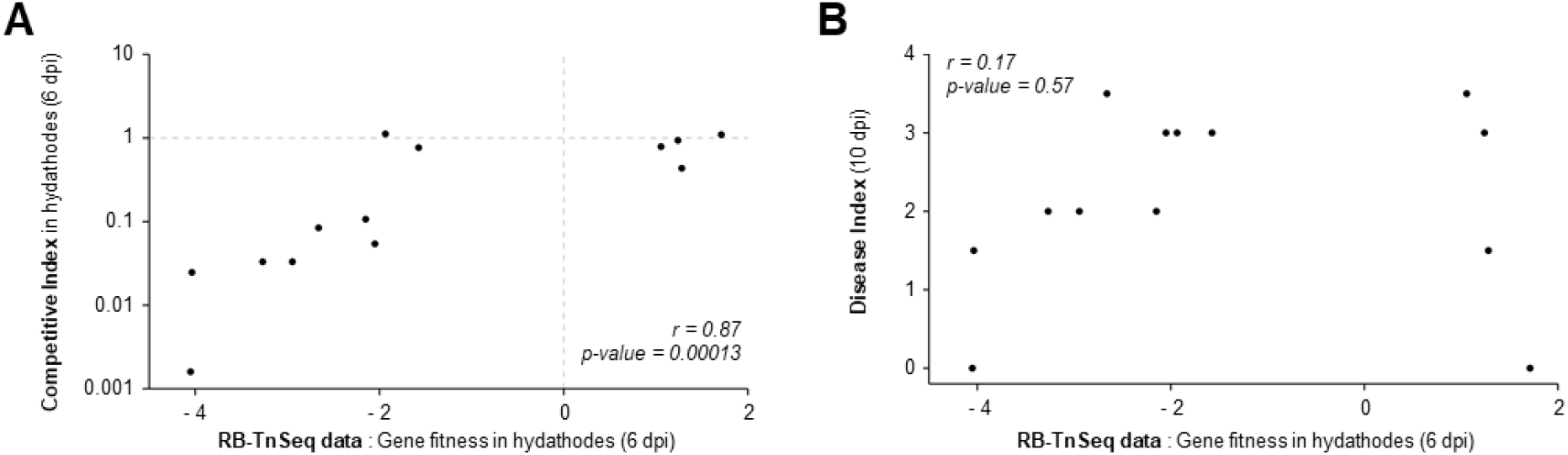
The importance of candidate genes identified by RB-TnSeq for fitness during plant infection is recapitulated by competition assays but is not well correlated with their contribution to disease severity. **A** Correlation between gene fitness values obtained with RB-TnSeq and 1:1 competition assays, both at 6 dpi in hydathodes. Dots represent the mean of gene fitness values for the 4 biological replicates of RB-TnSeq and the median of CI obtained over at least 4 biological replicates. **B** Correlation between gene fitness values obtained with RB-TnSeq at 6 dpi in hydathodes and disease symptoms scoring at 10 dpi. Dots represent the mean of gene fitness values for the 4 biological replicates of RB-TnSeq and the median of Disease Index obtained over 3 biological replicates. Correlation strength is indicated by the Spearman correlation coefficient *r* and the associated p-value.

**Fig. S5:**
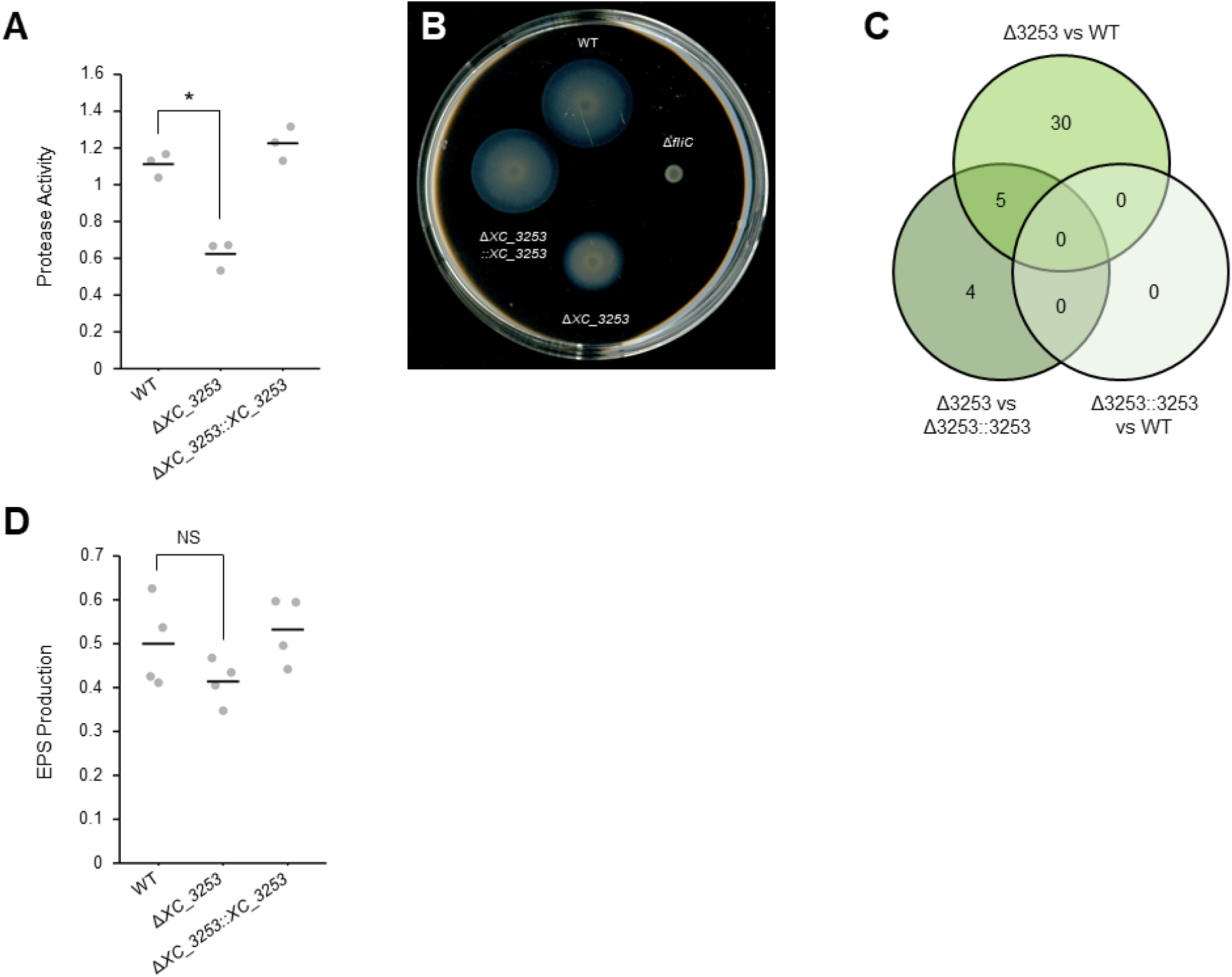
Impact of *XC_3253* on *Xcc* physiology. **A** Qualitative assay for total protease activity on skim milk agar plates. Activity for three independent replicates is given as an arbitrary unit calculated using this formula : PrA = [π(rhalo)^2^-π(rcolo)^2^]/ [π(rcolo)^2^] with “r” corresponding to the radius measured for colonies and halos of degradation. **B** Motility assay showing swimming abilities after 48h on soft agar plates. The WT strain and the Δ*fliC* non-motile flagellin mutant are positive and negative control of motility, respectively. **C** Differentially-expressed genes in the Δ*XC_3253* mutant and Δ*XC_3253::XC_33253* complemented strains. Genes were considered differentially-expressed if |log_2_FC| ≥ 1.5 and FDR ≤ 0.05. **D** Evaluation of exopolysaccharide production. Statistical significance of differences in (**A**) and (**D**) was assessed with the Wilcoxon test (*: p-value ≤ 0.05). Horizontal bars show the mean of at least 3 biological replicates.

**Fig. S6:**
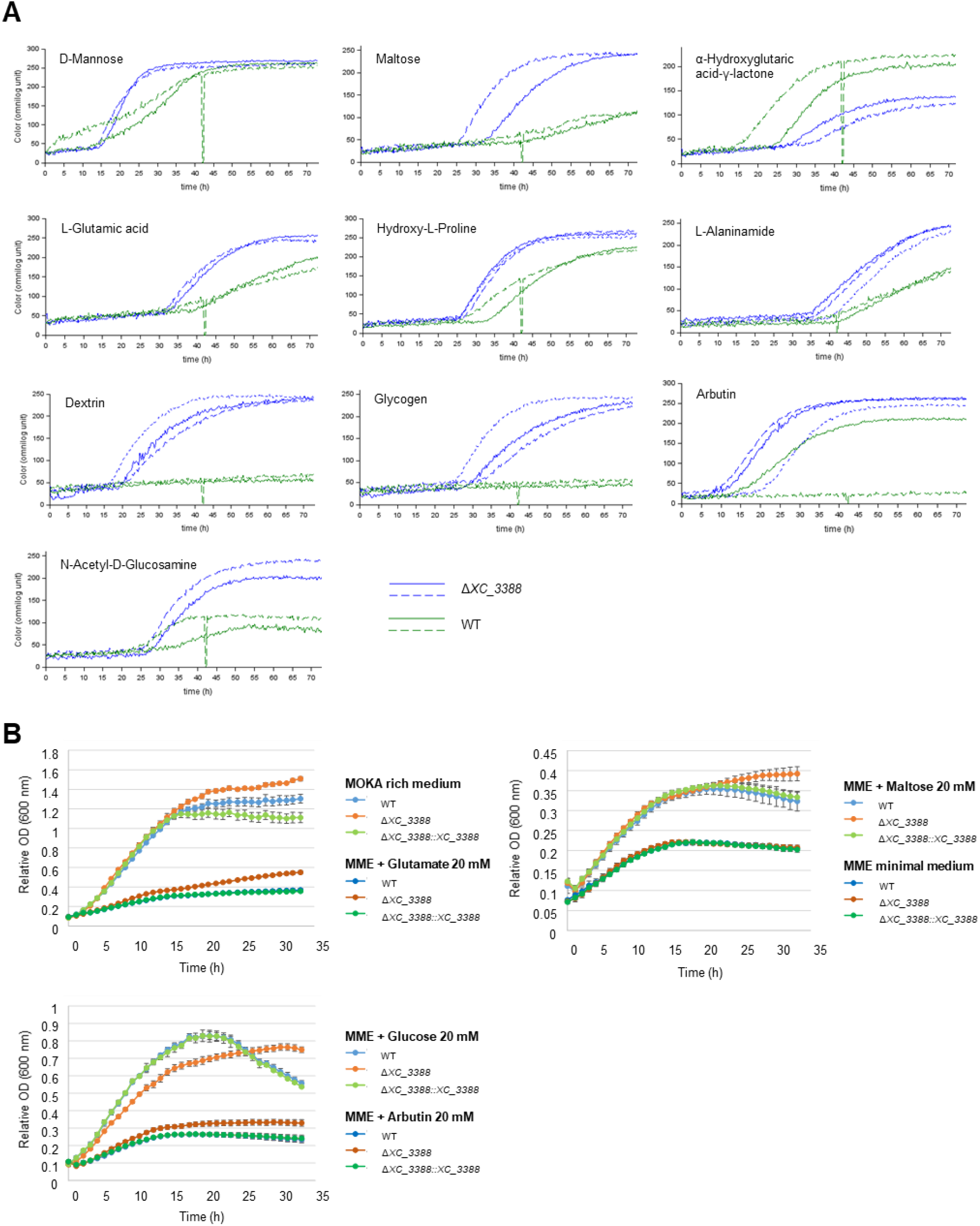
Impact of *XC_3388* on *Xcc* metabolism. **A** Metabolic fingerprinting on Biolog plates. The few carbon and nitrogen sources differentially metabolized between the Δ*XC_3388* (blue curves) and WT (green curves) strains are shown here. Two biological repetitions were performed except for the assay of Δ*XC_3388* on nitrogen source (plate PM03) for which a third repetition was required because of some variability. **B** Growth curves on diverse carbon sources confirm that *XC_3388* plays a role in *Xcc* multiplication, although Δ*XC_3388* shows different growth dynamics depending on the condition.

## References

Alexa A, Rahnenführer J, Lengauer T (2006) Improved scoring of functional groups from gene expression data by decorrelating GO graph structure. Bioinformatics 22: 1600–1607

An S-Q, Potnis N, Dow M, Vorhölter F-J, He Y-Q, Becker A, Teper D, Li Y, Wang N, Bleris L, et al (2020) Mechanistic insights into host adaptation, virulence and epidemiology of the phytopathogen Xanthomonas. FEMS Microbiol Rev 44: 1–32

Arlat M, Gough C, Barber CE, Boucher C, Daniels M (1991) Xanthomonas campestris Contains a Cluster of hrp Genes Related to the Larger hrp Cluster of Pseudomonas solanacearum. Mol Plant Microbe Interact 4: 593

Baek M, DiMaio F, Anishchenko I, Dauparas J, Ovchinnikov S, Lee GR, Wang J, Cong Q, Kinch LN, Schaeffer RD, et al (2021) Accurate prediction of protein structures and interactions using a three-track neural network. Science. doi: 10.1126/science.abj8754

Bai K, Yan H, Chen X, Lyu Q, Jiang N, Li J, Luo L (2022) The Role of RelA and SpoT on ppGpp Production, Stress Response, Growth Regulation, and Pathogenicity in Xanthomonas campestris pv. campestris. 9: 16

Blanvillain S, Meyer D, Boulanger A, Lautier M, Guynet C, Denancé N, Vasse J, Lauber E, Arlat M (2007) Plant Carbohydrate Scavenging through TonB-Dependent Receptors: A Feature Shared by Phytopathogenic and Aquatic Bacteria. PLOS ONE 2: e224

Büttner D, Bonas U (2010) Regulation and secretion of Xanthomonas virulence factors. FEMS Microbiol Rev 34: 107–133

Cain AK, Barquist L, Goodman AL, Paulsen IT, Parkhill J, van Opijnen T (2020) A decade of advances in transposon-insertion sequencing. Nat Rev Genet 1–15

Cerutti A, Jauneau A, Auriac M-C, Lauber E, Martinez Y, Chiarenza S, Leonhardt N, Berthomé R, Noel LD (2017) Immunity at cauliflower hydathodes controls infection by Xanthomonas campestris pv. campestris. Plant Physiol pp.01852.2016

Cerutti A, Jauneau A, Laufs P, Leonhardt N, Schattat MH, Berthomé R, Routaboul J-M, Noël LD (2019) Mangroves in the Leaves: Anatomy, Physiology, and Immunity of Epithemal Hydathodes. Annu Rev Phytopathol 57: 91–116

Diard M, Garcia V, Maier L, Remus-Emsermann MNP, Regoes RR, Ackermann M, Hardt W-D (2013) Stabilization of cooperative virulence by the expression of an avirulent phenotype. Nature 494: 353–356

Ditta G, Stanfield S, Corbin D, Helinski DR (1980) Broad host range DNA cloning system for gram-negative bacteria: construction of a gene bank of Rhizobium meliloti. Proc Natl Acad Sci 77: 7347–7351

Dugé de Bernonville T, Noël LD, SanCristobal M, Danoun S, Becker A, Soreau P, Arlat M, Lauber E (2014) Transcriptional reprogramming and phenotypical changes associated with growth of Xanthomonas campestris pv. campestris in cabbage xylem sap. FEMS Microbiol Ecol 89: 527–541

Duong DA, Jensen RV, Stevens AM (2018) Discovery of Pantoea stewartii ssp. stewartii genes important for survival in corn xylem through a Tn-Seq analysis. Mol Plant Pathol 19: 1929–1941

Figurski DH, Helinski DR (1979) Replication of an origin-containing derivative of plasmid RK2 dependent on a plasmid function provided in trans. Proc Natl Acad Sci 76: 1648–1652

Gerlin L, Cottret L, Cesbron S, Taghouti G, Jacques M-A, Genin S, Baroukh C (2020) Genome-Scale Investigation of the Metabolic Determinants Generating Bacterial Fastidious Growth. mSystems. doi: 10.1128/mSystems.00698-19

Gonzalez-Mula A, Lachat J, Mathias L, Naquin D, Lamouche F, Mergaert P, Faure D (2019) The biotroph Agrobacterium tumefaciens thrives in tumors by exploiting a wide spectrum of plant host metabolites. New Phytol 222: 455–467

He Y-Q, Zhang L, Jiang B-L, Zhang Z-C, Xu R-Q, Tang D-J, Qin J, Jiang W, Zhang X, Liao J, et al (2007a) Comparative and functional genomics reveals genetic diversity and determinants of host specificity among reference strains and a large collection of Chinese isolates of the phytopathogen Xanthomonas campestris pv. campestris. Genome Biol 8: R218

He Y-W, Boon C, Zhou L, Zhang L-H (2009) Co-regulation of Xanthomonas campestris virulence by quorum sensing and a novel two-component regulatory system RavS/RavR. Mol Microbiol 71: 1464–1476

He Y-W, Ng AY-J, Xu M, Lin K, Wang L-H, Dong Y-H, Zhang L-H (2007b) Xanthomonas campestris cell–cell communication involves a putative nucleotide receptor protein Clp and a hierarchical signalling network. Mol Microbiol 64: 281–292

Helmann TC, Deutschbauer AM, Lindow SE (2019) Genome-wide identification of Pseudomonas syringae genes required for fitness during colonization of the leaf surface and apoplast. Proc Natl Acad Sci 201908858

Helmann TC, Deutschbauer AM, Lindow SE (2020) Distinctiveness of genes contributing to growth of Pseudomonas syringae in diverse host plant species. PLOS ONE 15: e0239998

Irving SE, Choudhury NR, Corrigan RM (2021) The stringent response and physiological roles of (pp)pGpp in bacteria. Nat Rev Microbiol 19: 256–271

Jumper J, Evans R, Pritzel A, Green T, Figurnov M, Ronneberger O, Tunyasuvunakool K, Bates R, Žídek A, Potapenko A, et al (2021) Highly accurate protein structure prediction with AlphaFold. Nature 596: 583–589

Kim H, Joe A, Lee M, Yang S, Ma X, Ronald PC, Lee I (2019) A Genome-Scale Co-Functional Network of Xanthomonas Genes Can Accurately Reconstruct Regulatory Circuits Controlled by Two-Component Signaling Systems. Mol Cells. doi: 10.14348/molcells.2018.0403

Luneau JS, Cerutti A, Roux B, Carrère S, Jardinaud M-F, Gaillac A, Gris C, Lauber E, Berthomé R, Arlat M, et al (2022a) Xanthomonas transcriptome inside cauliflower hydathodes reveals bacterial virulence strategies and physiological adaptations at early infection stages. Mol Plant Pathol 23: 159–174

Luneau JS, Noël LD, Lauber E, Boulanger A (2022b) A β-glucuronidase (GUS) Based Bacterial Competition Assay to Assess Fine Differences in Fitness during Plant Infection. BIO-Protoc.

Macho AP, Zumaquero A, Ortiz-Martín I, Beuzón CR (2007) Competitive index in mixed infections: a sensitive and accurate assay for the genetic analysis of Pseudomonas syringae–plant interactions. Mol Plant Pathol 8: 437–450

Merda D, Briand M, Bosis E, Rousseau C, Portier P, Barret M, Jacques M-A, Fischer-Le Saux M (2017) Ancestral acquisitions, gene flow and multiple evolutionary trajectories of the type three secretion system and effectors in Xanthomonas plant pathogens. Mol Ecol 26: 5939–5952

Morris JJ, Lenski RE, Zinser ER (2012) The Black Queen Hypothesis: Evolution of Dependencies through Adaptive Gene Loss. mBio. doi: 10.1128/mBio.00036-12

van Opijnen T, Levin HL (2020) Transposon Insertion Sequencing, a Global Measure of Gene Function. Annu Rev Genet 54: null

Pandey SS, Patnana PK, Lomada SK, Tomar A, Chatterjee S (2016) Co-regulation of Iron Metabolism and Virulence Associated Functions by Iron and XibR, a Novel Iron Binding Transcription Factor, in the Plant Pathogen Xanthomonas. PLOS Pathog 12: e1006019

Price MN, Wetmore KM, Waters RJ, Callaghan M, Ray J, Liu H, Kuehl JV, Melnyk RA, Lamson JS, Suh Y, et al (2018) Mutant phenotypes for thousands of bacterial genes of unknown function. Nature 557: 503–509

Qian W, Han Z-J, He C (2008) Two-Component Signal Transduction Systems of Xanthomonas spp.: A Lesson from Genomics. Mol Plant Microbe Interact 21: 151–161

Qian W, Jia Y, Ren S-X, He Y-Q, Feng J-X, Lu L-F, Sun Q, Ying G, Tang D-J, Tang H, et al (2005) Comparative and functional genomic analyses of the pathogenicity of phytopathogen Xanthomonas campestris pv. campestris. Genome Res 15: 757–767

Robeson DJ, Bretschneider KE, Gonella MP (1989) A hydathode inoculation technique for the simulation of natural black rot infection of cabbage by Xanthomonas campestris pv. campestris. Ann Appl Biol 115: 455–459

Royet K, Parisot N, Rodrigue A, Gueguen E, Condemine G (2019) Identification by Tn-seq of Dickeya dadantii genes required for survival in chicory plants. Mol Plant Pathol 20: 287–306

Rufián JS, Macho AP, Corry DS, Mansfield JW, Ruiz-Albert J, Arnold DL, Beuzón CR (2018) Confocal microscopy reveals in planta dynamic interactions between pathogenic, avirulent and non-pathogenic Pseudomonas syringae strains. Mol Plant Pathol 19: 537–551

Rufián JS, Sánchez-Romero M-A, López-Márquez D, Macho AP, Mansfield JW, Arnold DL, Ruiz-Albert J, Casadesús J, Beuzón CR (2016) Pseudomonas syringae Differentiates into Phenotypically Distinct Subpopulations During Colonization of a Plant Host. Environ Microbiol 18: 3593–3605

Ryan RP, Fouhy Y, Lucey JF, Jiang B-L, He Y-Q, Feng J-X, Tang J-L, Dow JM (2007) Cyclic di-GMP signalling in the virulence and environmental adaptation of Xanthomonas campestris. Mol Microbiol 63: 429–442

Schäfer A, Tauch A, Jäger W, Kalinowski J, Thierbach G, Pühler A (1994) Small mobilizable multi-purpose cloning vectors derived from the Escherichia coli plasmids pK18 and pK19: selection of defined deletions in the chromosome of Corynebacterium glutamicum. Gene 145: 69–73

Sturm A, Heinemann M, Arnoldini M, Benecke A, Ackermann M, Benz M, Dormann J, Hardt W-D (2011) The Cost of Virulence: Retarded Growth of Salmonella Typhimurium Cells Expressing Type III Secretion System 1. PLOS Pathog 7: e1002143

Tang J, Tang D-J, Dubrow ZE, Bogdanove A, An S (2020) Xanthomonas campestris Pathovars. Trends Microbiol. doi: 10.1016/j.tim.2020.06.003

Tao J, He C (2010) Response regulator, VemR, positively regulates the virulence and adaptation of Xanthomonas campestris pv. campestris: VemR-regulated virulence and adaptation in Xcc. FEMS Microbiol Lett 304: 20–28

Taylor RK, Miller VL, Furlong DB, Mekalanos JJ (1987) Use of phoA gene fusions to identify a pilus colonization factor coordinately regulated with cholera toxin. Proc Natl Acad Sci 84: 2833–2837

Timilsina S, Potnis N, Newberry EA, Liyanapathiranage P, Iruegas-Bocardo F, White FF, Goss EM, Jones JB (2020) Xanthomonas diversity, virulence and plant–pathogen interactions. Nat Rev Microbiol. doi: 10.1038/s41579-020-0361-8

Torres M, Jiquel A, Jeanne E, Naquin D, Dessaux Y, Faure D (2022) Agrobacterium tumefaciens fitness genes involved in the colonization of plant tumors and roots. New Phytol 233: 905–918

Turner P, Barber C, Daniels M (1984) Behaviour of the transposons Tn5 and Tn7 in Xanthomonas campestris pv. campestris. Mol Gen Genet MGG 195: 101–107

Vicente JG, Holub EB (2013) Xanthomonas campestris pv. campestris (cause of black rot of crucifers) in the genomic era is still a worldwide threat to brassica crops. Mol Plant Pathol 14: 2–18

Vorholt JA (2012) Microbial life in the phyllosphere. Nat Rev Microbiol 10: 828–840

Wei K, Tang D-J, He Y-Q, Feng J-X, Jiang B-L, Lu G-T, Chen B, Tang J-L (2007) hpaR, a Putative marR Family Transcriptional Regulator, Is Positively Controlled by HrpG and HrpX and Involved in the Pathogenesis, Hypersensitive Response, and Extracellular Protease Production of Xanthomonas campestris Pathovar campestris. J Bacteriol 189: 2055–2062

Wetmore KM, Price MN, Waters RJ, Lamson JS, He J, Hoover CA, Blow MJ, Bristow J, Butland G, Arkin AP, et al (2015) Rapid quantification of mutant fitness in diverse bacteria by sequencing randomly bar-coded transposons. mBio 6: e00306–00315

